# Redox imbalance dictates dependence on GOT1 versus GOT2 for rod photoreceptor health during aging and stress

**DOI:** 10.64898/2026.04.05.716322

**Authors:** Meini Chen, Eric Weh, Moloy T. Goswami, Katherine M. Weh, Heather Hager, Peter Sajjakulnukit, Avi Weingarten, Shubha Subramanya, Nicholas Miller, Sraboni Chaudhury, Emma Piraino, Navdeep S. Chandel, Renee C. Ryals, Costas A. Lyssiotis, Thomas J. Wubben

## Abstract

Photoreceptor (PR) loss causes vision loss in many blinding diseases, and effective therapies to prevent this cell loss are lacking. Aspartate aminotransferases (GOTs), located in the cytosol (GOT1) and mitochondria (GOT2), are key components of the malate-aspartate shuttle, which transfers reducing equivalents from cytosol to mitochondria. Previous work has implicated the GOTs as potential modulators of blinding retinal disease. To determine the roles of GOT1 and GOT2 in rod PRs, we generated rod PR-specific *Got1* or *Got2* conditional knockout mice (*Got1* or *Got2* cKO). We previously showed that *Got1* cKO causes PR degeneration and is accompanied by NADH accumulation and a decreased retinal NAD^+^/NADH ratio. Here, we show that NADH oxidation via metabolic or genetic means prolongs PR survival in *Got1* cKO animals, implicating NADH accumulation, or reductive stress, as a key driver of PR degeneration. In contrast, *Got2* cKO causes minimal PR degeneration and alterations in retinal NADH and the NAD^+^/NADH ratio that oppose reductive stress. Interestingly, GOT2, but not GOT1, is decreased in multiple models of PR degeneration, including retinal detachment (RD) where the NAD^+^/NADH ratio favors a reductive state. Notably, loss of *Got2* in PRs demonstrates a neuroprotective effect after experimental RD suggesting decreased GOT2 expression may be part of a stress response to promote PR survival. Overall, this study illustrates the differential dependence on the GOTs for PR health, provides evidence that an overly reductive environment is detrimental to PR survival, and identifies GOT2 as a novel therapeutic target with potentially broad application in blinding diseases.

## Introduction

Photoreceptor (PR) cell death is the ultimate cause of vison loss in many acquired and inherited retinal diseases (IRDs), such as retinal detachment, retinitis pigmentosa (RP), and age-related macular degeneration (AMD) [1–3]. Greater than 200 million people suffer from these blinding diseases worldwide and this number is projected to increase as the population ages [4]. Gene therapy for IRDs offers hope, but given the vast genetic heterogeneity and the enormous complexity of clinical development, regulatory approval of precision medicine strategies for each IRD is not feasible [3]. Current therapies for patients with AMD are either effective for only a small percentage of patients with the exudative form (i.e. anti-VEGF therapy) or demonstrate modest clinical benefits in the late-stage, nonexudative form (i.e. complement inhibitors). Neither of these therapeutic strategies prevent vision loss over time [5,6]. As such, millions with retinal degenerative diseases remain without effective therapies to prevent vision loss, so a significant and urgent unmet need exists to identify novel neuroprotective approaches with broad therapeutic potential to prevent vision loss in these blinding diseases. Targeting molecular pathways fundamental to PR degeneration is a potential solution to this unmet need by offering more widely implementable, innovative therapeutic strategies.

PRs are one of the most metabolically active cells in the body due to both the massive energy demands of phototransduction and the continuous biosynthetic requirements for outer segment renewal. At the same time, PRs have limited reserve capacity to meet these metabolic demands, so they are uniquely susceptible to small changes in metabolic homeostasis. This is supported by the occurrence of isolated retinal degeneration phenotypes in humans caused by variants in genes encoding for ubiquitously expressed metabolic enzymes [3,7–9]. Therefore, maintaining metabolic homeostasis is critical to PR survival.

Accumulating evidence indicates that targeting PR metabolism to increase PR resistance to stress is an attractive neuroprotective strategy. For example, PRs are metabolically dependent on glucose to generate energy and anabolic building blocks for baseline physiologic functions, and disruption of glucose metabolism is a key driver of PR degeneration [10–14]. Multiple preclinical strategies to boost glycolysis are being investigated and a current Phase 1/2 trial is evaluating subretinal gene therapy to stimulate PR glucose metabolism for IRD treatment (NCT05748873) [15–18]. However, beyond glucose metabolism and glycolysis, the metabolic pathways and adaptations integral to promoting PR health at baseline and under stress remain largely unknown. This is a critical gap in our knowledge as identification of these pathways is expected to reveal new therapeutic targets.

The nicotinamide adenine dinucleotide (NAD^+^)/reduced NAD^+^ (NADH) redox couple and its interconversion is essential for PRs to meet their immense metabolic demands. NADH donates electrons to fuel mitochondrial energy production, whereas the regeneration of NAD^+^ is necessary to maintain glycolysis and other cytosolic oxidative pathways [19–21]. Because the inner mitochondrial membrane is impermeable to NADH, reducing equivalents are transported into the mitochondrial matrix indirectly via the malate-aspartate shuttle (MAS) or the glycerol 3-phosphate shuttle [22,23]. The MAS is the major metabolic redox shuttle in PRs, as its components are highly expressed and their activity is particularly enriched in PR inner segments [24,25]. Conversely, the glycerol-3-phosphate shuttle plays a limited role due to the low abundance of transcripts (GSE169047 and 63473) and likely absence of glycerol-3-phosphate dehydrogenase in PRs [26–29].

Previous work has demonstrated that the MAS is important for both retinal function and metabolism [19,24,30]. Glutamic-oxaloacetic transaminase (GOT), also known as aspartate aminotransferase, exists as two isozymes in mammals. These isozymes vary in sub-cellular location, either cytoplasmic (GOT1) or mitochondrial (GOT2), and are crucial components of the MAS [23]. GOT1 catalyzes the transamination of aspartate (Asp) and α-ketoglutarate (α-KG) to generate oxaloacetate (OAA) and glutamate (Glu) in the cytoplasm, whereas GOT2 catalyzes the reverse reaction in the mitochondrial matrix [23]. The metabolites interconverted by the GOTs serve as pivotal intermediates in the TCA cycle, amino acid biosynthesis, and glutamine metabolism [23,24,31–33]. Coupling this with the GOTs’ well-known role in the MAS highlights their essential role in coordinating cellular energy production, redox balance, and biosynthetic processes. Given these critical metabolic roles, there has been increased interest in the function of GOT1 and GOT2 in the retina. To this end, the retina has been shown to rely on aspartate aminotransferases for amino acid metabolism *ex vivo* [34], and the expression of both GOT1 and GOT2 is altered early in the retinal degeneration of a preclinical model of RP [35], suggesting their therapeutic potential as PR neuroprotective targets.

To advance our understanding of the importance of the MAS, and particularly GOT1 and GOT2, in PRs *in vivo* and potentially identify therapeutic targets that promote PR resistance to stress, we generated mouse models that lacked either *Got1* or *Got2* specifically in rod PRs. Previous work from our lab demonstrated that *Got1* deficiency in rods results in PR degeneration [24]. Here, we show that this PR degeneration is driven by NADH accumulation, or reductive stress. Furthermore, in contrast to *Got1*, the loss of *Got2* induces minimal PR degeneration with a distinct metabolic profile. Excitingly, we found that GOT2 suppression in rod PRs is neuroprotective in an experimental model of retinal detachment. Taken with our data demonstrating that GOT2 expression is decreased in numerous preclinical models of PR degeneration, our study suggests that targeting GOT2 may be a viable therapeutic strategy to prevent PR cell death in disease.

## Materials and Methods

### Animals

All animal experiments were performed with permission from the Institutional Animal Care & Use Committee at the University of Michigan (Protocol number: PRO00012747 or PRO00012733) and in compliance with the Association for Research in Vision and Ophthalmology Statement for the Use of Animals in Ophthalmic and Vision Research. Mice were housed at room temperature (RT) in 12-hour light/12-hour dark cycles with free access to food and water unless described otherwise. To ensure the retinal phenotype was secondary to the loss of the gene of interest, mice were confirmed to lack the *rd8* mutation [36]. The mice utilized in this study, which are on the C57BL/6J background, were also confirmed to carry the loss-of-function nicotinamide nucleotide transhydrogenase (*Nnt*) mutation [37]. Both male and female mice were used for all experiments. Transgenic animals with conditional deletion of *Got1* from rod PRs (*Got1^fl/fl^*;*Rho-Cre^+^*, *Got1* cKO) were created by crossing animals carrying Lox-P sites flanking exon 3 of the *Got1* gene to mice harboring a Cre-recombinase under the control of the rhodopsin promoter [24,38]. Mice carrying Lox-P sites flanking exon 2 of the *Got2* gene were a generous gift from Dr. Costas Lyssiotis and originally generated by Ozgene [32]. Animals with conditional deletion of *Got2* from rod PRs (*Got2^fl/fl^;Rho-Cre+*, *Got2* cKO) were produced using a breeding scheme similar to that above for *Got1* cKO. Animals carrying only the Cre-recombinase without the *Got1* or *Got2* floxed alleles were used as controls (*Got1^wt/w^*^t^*;Rho-Cre+*, *Got1* WT; *Got2^wt/wt^;Rho-Cre+*, *Got2* WT). *CytoLbNOX^LSL/LSL^* mice carrying a Lox-P-Stop-Lox-P (LSL) cassette upstream of *cytoLbNOX*, followed by an internal ribosome entry site (IRES)-linked enhanced green fluorescent protein (eGFP), were generously provided by Dr. Navdeep Chandel at Northwestern University [39]. To achieve rod PR-conditional expression of cytoLbNOX, *cytoLbNOX^LSL/LSL^* mice were crossed with *Got1* cKO mice or *Got1* WT mice, generating *cytoLbNox/Got1* cKO mice (*cytoLbNOX^LSL/LSL^;Got1^fl/fl^*;*Rho-Cre^+^*) and *cytoLbNox/Got1* WT (*cytoLbNOX^LSL/LSL^;Got1^wt/wt^*;*Rho-Cre^+^*) mice.

*Rd2* and *rd12* mice were bred and maintained at Oregon Health & Science University. Mice were housed in standard conditions under a 12/12-hour light-dark cycle. All experiments were approved by the Institutional Animal Care and Use Committee at OHSU (IACUC Protocol TR03_IP00000610), and adhered to the ARVO Statement for the Use of Animals in Ophthalmic and Vision Research.

### Rescue studies

For rescue experiments, MCTI-566 (25 mg/kg) [40] or vehicle (40% 2-hydroxypropyl-β-cyclodextrin (Cayman Chemical, Ann Arbor, MI; Cat #16169 in PBS)) was injected via the intraperitoneal (IP) route into *Got1* cKO mice once every other week starting at 2 months of age. In separate experiments, sodium pyruvate (Millipore-Sigma, St. Louis, MO, USA; Cat #P5280) was added to the drinking water (2 mg/mL) of *Got1* cKO mice starting at 2 months of age and refreshed weekly [41].

### Immunofluorescence

For paraffin sectioning, mouse eyes were enucleated, fixed overnight in 10% neutral buffered formalin (Epredia, Netherlands B.V.; Cat #511201), embedded in paraffin and sliced into 4 µm thick sections. Deparaffinization of the sections and antigen retrieval using citrate buffer (pH 6.0) was carried out as previously published [42]. Sections were blocked using 10% normal goat serum (NGS, MilliporeSigma, Cat #G9023) or normal donkey serum (NDS, MilliporeSigma, Cat #D9663), supplemented with 1% bovine serum albumin (BSA, Millipore-Sigma, Cat #A9647) in PBST (1X PBS with 0.125% Triton X-100) for 1 hour at RT. Blocking solution was replaced with primary antibody diluted in wash solution (1% BSA and 1% NGS or NDS in PBST) and sections were placed in a humidified chamber overnight at 4°C. The following day, sections were washed three times with wash solution before incubation with secondary antibody (1 hour, RT, light protected). Sections were rinsed with PBS three times. A 30 μL drop of ProLong Gold Antifade with DAPI (Thermo Fisher Scientific, Waltham, MA, USA; Cat #P36935) was applied to the slides and covered with a coverslip.

For cryosections, mouse eyes were enucleated, fixed in 4% paraformaldehyde (Electron Microscopy Sciences, Hatfield, PA, USA; Cat #15713) in PBS for 2 hours at 4°C. Retinas were dissected in PBS, cryo-preserved in 30% sucrose (MilliporeSigma, Cat #S0389) overnight at 4°C, embedded in Optimal Cutting Temperature (O.C.T) compound (Thermo Fisher Scientific, Cat #4585), cryosectioned at 10 µm, and stored at −20°C. Sections were dried in a 37°C incubator for 30 min, and washed three times for 5 min each with PBST (1X PBS with 0.1% Triton X-100) at RT. Sections were blocked using blocking buffer (10% horse serum (Thermo Fisher Scientific, Cat #26050070) supplemented with 0.4% Triton X-100 in PBS) for two hours at RT and then incubated with primary antibodies diluted in blocking buffer overnight at 4°C. The following day sections were washed three times with PBST and incubated with secondary antibodies for 2 hr at RT. After washing three times in PBST, sections were mounted with ProLong Gold Antifade Mountant with DAPI (Thermo Fisher Scientific) and covered with a coverslip [43].

Images were captured using a Leica DM6000 microscope (Leica Microsystems, Wetzlar, Germany) equipped with a 40X objective and acquired using LAS X Office software (Leica Microsystems). The antibodies utilized in this study are listed in Table 1.

**Table 1.** List of antibodies utilized in this study.

| Antibody | Source | Dilution | Manufacturer (Cat #) |
| --- | --- | --- | --- |
| GOT1 | Mouse | 1:1000 (WB) | Abcam (ab239487) |
|  |  | 1:200 (IF) |  |
| GOT2 | Rabbit | 1:1000 (WB) | Atlas antibodies (HPA018139) |
| FABP1 (GOT2) | Rabbit | 1:300 (IF) | Abcam (ab153924) |
| HSP90 | Rabbit | 1:1000 (WB) | Cell Signaling Technology<br>(4877S) |
| OPN1MW | Goat | 1:200 (IF) | Santa Cruz (sc22117) |
| Total OXPHOS | Mouse | 1:1000 (WB) | Abcam (ab110413) |
| TIM23 | Mouse | 1:1000 (WB) | BD Biosciences (611223) |
| eGFP | Mouse | 1:4000 (WB) | TAKARA (632381) |
| eGFP | Chicken | 1:400 (IF) | Invitrogen (A10262) |
| Alexa Fluor 488<br>Anti-rabbit | Donkey | 1:500 (IF) | Jackson ImmunoResearch<br>Laboratories (711-545-152) |
| Alexa Fluor 594<br>Anti-mouse | Donkey | 1:400 (IF) | Jackson ImmunoResearch<br>Laboratories (715-585-151) |
| Alexa Fluor 647<br>Anti-goat | Donkey | 1:2000 (IF) | Invitrogen (A21447) |
| Alexa Fluor 488<br>Anti-chicken | Donkey | 1:400 (IF) | Sigma (SAB4600031) |
| Anti-rabbit IgG | Goat | 1:2500 (WB) | Cell Signaling Technology<br>(7074S) |
| Anti-mouse IgG | Horse | 1:2500 (WB) | Cell Signaling Technology<br>(7076S) |

### Histology and image analysis

Mouse eyes were harvested and fixed in 10% neutral buffered formalin (Epredia), embedded in paraffin, and sectioned to 4 µm thickness. Retinal sections through the optic nerve were selected and deparaffinized as previously described [42]. Sections were stained with hematoxylin and eosin following our standard protocol [24]. Retinal images were acquired using a Leica DM6000 microscope with a 20X objective and acquired using LAS X Office software.

### Western Blotting

Whole retinas were harvested and lysed in RIPA buffer (Thermo Fisher Scientific, Cat #89900) containing protease and phosphatase inhibitors (Cell Signaling Technology, Denver, MA, USA; Cat #5872S). Immunoblots were performed as previously described [42]. Briefly, samples were homogenized, centrifuged and protein concentrations were determined using the Pierce^TM^ BCA kit (Thermo Fisher Scientific, Cat #23225). Equivalent micrograms (µg) of protein from each sample were mixed with 4X Laemmli sample buffer (Bio-Rad, Cat #1610747) supplemented with β-mercaptoethanol (MilliporeSigma, Cat #M6250). Most samples were boiled for 5 minutes at 95°C; however, samples blotted with the total OXPHOS primary antibody cocktail (Table 1) were boiled at 50°C for 5 minutes following the manufacturer’s instructions. Proteins were separated using 4-20% precast polyacrylamide gels (Bio-Rad, Cat #5671094) and transferred to a PVDF membrane using the TurboBlot transfer system (Bio-Rad, Cat #1704150). After transfer, blots were incubated in 5% non-fat milk in TBS (Tris-buffered Saline, Bio-Rad, Cat #1706435), supplemented with 0.1% Tween-20 (Thermo Fisher Scientific, Cat #28320) for 1.5 hours at RT. Blots were incubated overnight at 4°C with primary antibody diluted in 5% BSA, except for the OXPHOS antibody cocktail, which was diluted in 1% milk according to the manufacturer’s instructions. The following day, membranes were rinsed with TBST before a 1 h RT incubation with secondary antibody (diluted in 5% non-fat milk). Blots were developed using chemiluminescence solution (EcoBright Femto HRP 100, Innovative Solutions, MI, USA; Cat #EBFH100) for 5 minutes before imaging with an Azure 600 imaging system (Azure Biosystems, Dublin, CA, USA). ImageJ was used to quantify the protein bands. Antibodies used for western blotting are shown in Table 1. Retinas from *rd2* and *rd12* mice were harvested at the Oregon Health & Science University, flash frozen, and shipped to the University of Michigan on dry ice. Once arrived, retinas were processed in a similar manner as described above.

### Quantitative real-time PCR (qRT-PCR)

Whole retinas were harvested and immediately preserved in RNAlater solution (Thermo Fisher Scientific, Cat #AM7020). Total RNA was extracted using the RNeasy Mini Kit (Qiagen, Cat #74104) following the manufacturer’s protocol prior to quantitation using a Nanodrop 1000 (Thermo Fisher Scientific). Approximately 500 ng of total RNA was used as input for cDNA synthesis using the RNA QuantiTect Reverse transcription kit (Qiagen, Cat #205311). For each qRT-PCR reaction, approximately 10 ng of cDNA was used as a template and combined with the PowerTrack SYBR Green Supermix (Applied Biosystems, Waltham, MA, USA; Cat #A46109) as previously described [24]. A CFX Opus 384 Real-Time PCR System (Bio-Rad) was used to perform qRT-PCR. *Actb* was used as the housekeeping gene to calculate relative gene expression levels using the 2^-ΔΔCt^ method [44]. Primers used in the present study are listed in Supplementary Table 1.

### Optical coherence tomography (OCT) and Electroretinography (ERG)

Animals received 1% tropicamide and 2.5% phenylephrine eye drops to dilate their pupils before being anesthetized with ketamine/xylazine at a dose of 90/10 mg/kg. OCT was performed using the Envisu-R SD-OCT system (Leica Microsystems Inc., Buffalo Grove, IL, USA) to capture frames from a 1.4 mm B-scan and a 1.4 mm × 1.4 mm rectangular volume scan as previously described [45]. The thickness of three retinal layers (total retina, combined inner segment/outer segment (IS/OS), and outer nuclear layer (ONL)) was measured with the Diver 1.0 software (Leica Microsystems) after first registering and averaging the image data. The average thickness of total retina, IS/OS, and ONL was measured at 16 points, spaced 140 µm apart, starting from the optic nerve head and following the 9×9 template [45].

For retinal function assessment, ERG using a Diagnosys Celeris ERG instrument (Diagnosys LLC, Lowell, MA, USA) was performed on mice that were dark-adapted overnight prior to measurements. Both scotopic and photopic responses were recorded and analyzed using Espion V6 software (Diagnosys LLC), as we previously published [46].

### Unlabeled targeted metabolomics

For unlabeled targeted metabolomics, both retinas of each animal were harvested, combined and snap-frozen on dry ice as previously described [46]. The tissue was weighed before adding ice-cold 80% methanol and homogenized using an OMNI Bead Ruptor (OMNI International, Kennesaw, GA, USA; Cat #19-050A) to extract metabolites. Lysates were centrifuged at 14,000 x g for 10 minutes at 4°C with the supernatant stored at −80°C until further processing. An additional sample was collected from each group, weighed and homogenized in RIPA buffer prior to protein quantitation using a BCA assay as described above. The protein concentration for each group (µg protein/mg tissue) was used to normalize the volume of supernatant for lyophilization of each sample using a SpeedVac concentrator (Thermo Fisher Scientific, Cat #13875355). Dried metabolite pellets were resuspended in a 1:1 methanol/water solution and analyzed by liquid chromatography-coupled tandem mass spectrometry (LC-MS/MS) using an Agilent Technologies Triple Quad 6470 system (Santa Clara, CA, USA) as previously described [46]. The targeted method consists of 215 analytes from the Agilent Metabolomics Dynamic MRM (dMRM) Database and Methods with additional analytes added to this method as recently described [47]. Agilent MassHunter Workstation Quantitative Analysis for QQQ (Version B.10.1.733.009.) was used to process raw data with additional statistical analyses conducted using Microsoft Excel. Each sample was normalized by the total intensity of all metabolites to scale equal sample loading. Fold change in metabolite abundance between control and experimental groups was calculated by normalizing each sample’s metabolite abundance to the mean abundance of all control samples.

### NAD^+^/NADH Measurements

The NAD^+^/NADH ratio was measured in attached and detached retinas from adult Brown Norway rats using the NAD^+^/NADH-Glo Assay (Promega, Madison, WI, USA, Cat #G9071) as previously described [46]. Briefly, whole retinas were harvested 24 h after experimental retinal detachment and homogenized in 300 µL PBS/bicarbonate buffer containing 0.5% dodecyl trimethylammonium bromide (DTAB). For NAD^+^ measurements, the lysate was mixed with 0.4 N HCl (Thermo Fisher Scientific, Cat #A144), heated to 60°C for 15 min, cooled to RT for 10 min and neutralized with 0.5 M Trizma base (Millipore-Sigma, Cat #T1503). For NADH measurements, the remaining lysate was heated to 60°C for 15 min, cooled to RT for 10 min and Trizma/HCl solution was added to the sample. Following lysate preparation, samples were incubated at RT for 30 min with equal amount of the NAD^+^/NADH-Glo detection reagent in a 96-well white walled tissue culture plate (Thermo Fisher Scientific, Cat #3610). Luminescence was recorded using the Omega plate reader (BMG Labtech) and data are presented as NAD^+^/NADH ratio.

### Retinal detachment

Experimental retinal detachment was created in mice and rats as previously described [48]. Briefly, eyes were dilated, and the animal anesthetized as noted above before a drop of 0.5% Proparacaine/HCl ophthalmic solution (Bausch & Lomb, Bridgewater, NJ, USA; Cat# 24208-730-06) was applied to the ocular surface. GenTeal Tears Lubricant Eye Gel (Alcon, Fort Worth, TX, USA) was placed on the cornea as a coupling agent for a 3 or 5 mm glass coverslip to facilitate visualization of the posterior segment. A sclerotomy was performed just posterior to the limbus using a 25-gauge (G) microvitreoretinal blade taking care to avoid damaging the lens. A 35G beveled needle attached to a NanoFil syringe (World Precision Instruments, Sarasota, FL, USA; Cat #s NF35BV-2, NANOFIL-100) was passed through the sclerotomy and tunneled underneath the neural retina via a retinotomy in the peripheral retina. A micropump was used to inject 4 µL of Healon (Abbott Medical Optics, Santa Ana, CA, USA, Cat #05047450842) into the subretinal space, resulting in detachment of at least one-half to two-thirds of the neural retina from the underlying retinal pigment epithelium (RPE). Animals were euthanized and retinas harvested one day, three days, or seven days later as described above. Eyes that underwent sclerotomy only served as the attached retina control group.

### Flow cytometry

Retinas were freshly harvested and processed for TUNEL staining as previously described [49]. In brief, each retina was digested with 700 µL of 0.25% Trypsin-EDTA (Gibco, Cat #25200-56) supplemented with 400 µg of deoxyribonuclease II (Worthington Biochemical, Lakewood, NJ, USA; Cat #LS002425) at RT for 20 min with gentle agitation. The tissue was repeatedly passed through an 18G needle before adding 2 mL of FACS buffer (1% FBS in PBS) to inactivate the trypsin solution. Resuspended cells were fixed with 1% paraformaldehyde diluted in PBS for 20 minutes at 4°C and permeabilized using 70% ethanol for at least 4 h. Samples were further permeabilized using 0.2% TritonX-100 (MilliporeSigma, Cat #T8787) in PBS for 5 min at RT. Cells were washed and resuspended before staining using the DeadEnd Fluorometric TUNEL kit (Promega, Cat #G3250). Stained samples were analyzed using an Attune NXT flow cytometer (Thermo Fisher Scientific; FITC channel (530 nm)). Collected data were analyzed with FlowJo v10.10.0 software (BD Biosciences, Franklin Lakes, NJ, USA). The gating strategy for analysis of flow samples was similar to that previously described (Supp. Fig. 1) [49].

### BaroFuse analysis

The BaroFuse (Entox Biosciences; Mercer Island, WA, USA) was used to measure oxygen consumption rate (OCR) in fresh *ex vivo* retinal tissue as previously described [46,48,50]. Each tissue chamber was loaded with a single, freshly harvested, whole retina. A chamber without tissue was used as a negative control. HEPES-buffered Krebs-Ringer Solution (KRB; Thermo Fisher, Cat #J67795.K2), supplemented with 0.1 g/100 mL BSA and glucose to achieve a final 5.5 mM glucose concentration (Millipore-Sigma, Cat #G8270) was added to the media reservoirs. The atmosphere in the media reservoirs was controlled to achieve a desired gas composition (21% oxygen, 5% CO2, 74% N2) within the media and the temperature was maintained at 37°C. Oligomycin-A (Cayman Chemical, Ann Arbor, MI, USA, Cat #11342), FCCP (Trifluoromethoxy carbonylcyanide phenylhydrazone, Cayman Chemical, Cat #15218) and KCN (Potassium Cyanide, Thermo Fisher, Cat #012136) were injected into the perifusion media at different time points via the injection port. Data was analyzed using BaroFuse Data Processor Software (Entox Biosciences).

### Statistical analysis

All data are presented as meanL±LSEM. Statistical analysis was performed using GraphPad Prism 10 software. Data were analyzed using Student’s t-test for two-group comparisons and one-way or two-way ANOVA for multiple comparisons, followed by Bonferroni’s post hoc test when applicable. PL<L0.05 was considered as statistically significant.

## Results

### *Got2* rod PR-specific conditional knockout causes minimal retinal degeneration

In order to evaluate the role of GOT1 and GOT2 in PR function and survival, we generated rod PR-specific, *Got1* or *Got2* conditional knockout (cKO) mice, respectively. The successful establishment of a rod PR-specific *Got1* cKO mouse was previously described by our group with the loss of GOT1 in rod PRs resulting in progressive, robust PR degeneration and functional loss [24]. Specifically, inner segment/outer segment (IS/OS) thinning was observed as early as 2 months of age followed by outer nuclear layer (ONL) thinning beginning at 4 months of age with concordant retinal function impairment [24].

To determine whether loss of GOT2 impacts PR survival and function in a similar manner, a rod PR-specific *Got2* cKO mouse was created and characterized here. Immunofluorescence on retinal sections from 2-month-old mice demonstrated loss of GOT2 expression in rod PR inner segments of the *Got2* cKO mouse with expression remaining in cone inner segments (white arrows, Fig. 1A). Western blot analysis further confirmed a significant reduction in GOT2 protein levels in total retinal lysate harvested from 2-month-old *Got2* cKO compared to *Got2* WT mice, without a compensatory upregulation of the cytosolic isoform GOT1 (Fig. 1B).

**Fig. 1.**
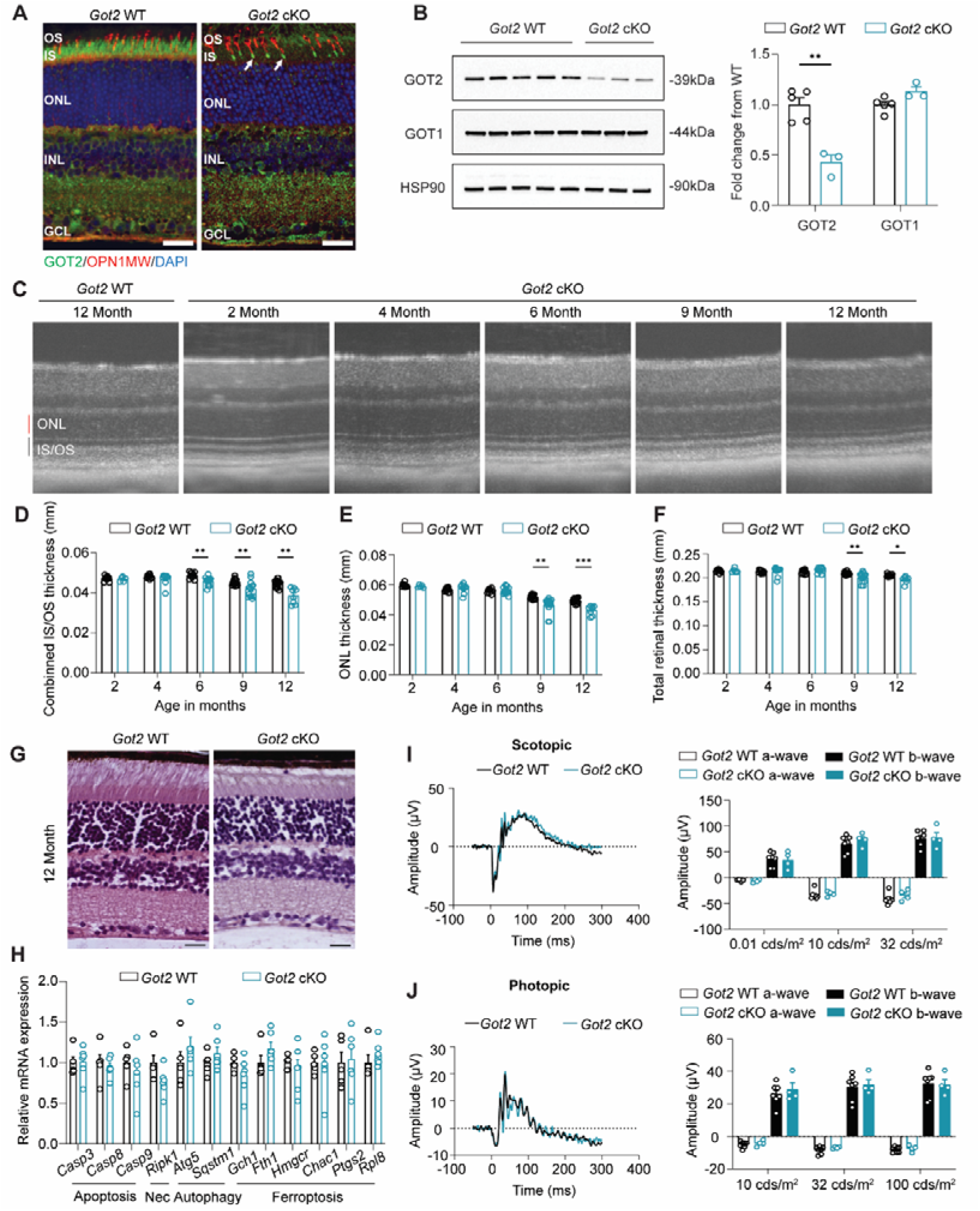
Rod-specific deletion of GOT2 leads to minimal PR degeneration. **(A)** Representative immunofluorescence images of 2-month-old retina stained for GOT2 (green), OPN1MW (red), and nuclei (DAPI) in *Got2* WT and *Got2* cKO mice; Scale bar: 25 µm; **(B)** Western blot analysis shows decreased GOT2 protein levels in the *Got2* cKO retina, with unchanged GOT1 protein levels; N=3-5 animals per group; **(C)** Representative OCT images show modest structural changes in *Got2* cKO compared to WT mice from 2 to 12 months; **(D-F)** Quantitation of inner segment/outer segment (IS/OS), outer nuclear layer (ONL), and total retinal thickness over time as determined by OCT; N=3-8 animals per group; **(G)** Histology shows modest structural changes in the *Got2* cKO mice at 12 months of age; Scale bar: 25 µm; **(H)** qRT-PCR of genes related to cell death pathways including apoptosis, necroptosis (Nec.), autophagy and ferroptosis shows no significant changes between *Got2* WT and cKO retina at 2 months of age; N=5-6 animals per group; Representative scotopic (32 cd·s/m²) **(I)** and photopic ERG tracings (100 cd·s/m²) **(J)** and quantification of respective a- and b-wave amplitudes in *Got2* cKO and WT mice at 12 months of age; N=4-7 animals per group; Statistical differences are based on an unpaired two-tail student’s t-test as compared to WT mice; *P <0.05, **P<0.01 and ***P<0.001. Mean ± SEM. HSP90: heat shock protein 90.

As the *Got2* cKO mouse successfully downregulated GOT2 in rod PRs, we next sought to determine the impact of this cKO on long-term PR survival. Retinal thickness was assessed using optical coherence tomography (OCT) at multiple time points from 2 to 12 months of age in *Got2* cKO and WT mice. Loss of GOT2 leads to a small, but statistically significant, progressive reduction in IS/OS thickness starting at 6 months of age. The ONL and total retinal thickness are preserved until 9 months of age where small, but statistically significant decreases become detectable (Fig. 1C-F).

Histology confirmed the minimal ONL changes in *Got2* cKO retinas at 12 months of age (Fig. 1G). qRT-PCR analysis of genes involved in the cell death pathways of apoptosis, necroptosis, ferroptosis or autophagy at 2 months of age, a time prior to ONL thinning, did not reveal any significant changes in expression (Fig. 1H), which is in contrast to the upregulation of apoptotic genes in the *Got1* cKO mouse at a similar age [24]. Moreover, even at 12 months of age, ERG analysis revealed no measurable differences in a- or b-wave amplitudes between *Got2* cKO and WT animals under either scotopic or photopic conditions (Fig. 1I & J). These data demonstrate that in contrast to GOT1, loss of GOT2 in rod PRs results in a much more modest retinal phenotype.

### *Got2* cKO impacts the retinal NAD⁺/NADH ratio differently than *Got1* cKO

We previously showed that loss of GOT1 in rod PRs causes a significant increase in retinal NADH with a corresponding decrease in the NAD^+^/NADH ratio [24]. Given that both GOT1 and GOT2 coordinate the transfer of reducing equivalents from the cytosol to the mitochondria (Fig. 2A), we sought to determine the impact of *Got2* cKO on the NAD^+^/NADH ratio. Targeted liquid chromatography-tandem mass spectrometry (LC-MS/MS) metabolomics was performed on retinas from 2-month-old *Got2* cKO and WT mice. In stark contrast to that observed in the *Got1* cKO retina, a significant decrease in NADH without any change in NAD^+^ was observed in the *Got2* cKO compared to WT retina, resulting in an increased NAD^+^/NADH ratio (Fig. 2B). Beyond the NAD^+^/NADH ratio, disruption of the MAS *in vitro* has been shown to impact the levels of metabolites involved in the MAS, glycolysis, TCA cycle, and serine biosynthesis [23,31,32,51,52]. However, metabolites involved in the MAS, including aspartate, malate, glutamate and α-ketoglutarate, did not show significant changes in *Got2* cKO compared to WT retina *in vivo*. Furthermore, only a few metabolites involved in glycolysis and the TCA cycle were altered in the *Got2* cKO retina, and serine levels were unchanged (Fig. 2C). Similarly, in our previous study, only a few glycolytic and no TCA cycle metabolites showed significant changes in the *Got1* cKO retina despite the significantly altered NAD^+^/NADH ratio [24].

**Fig. 2.**
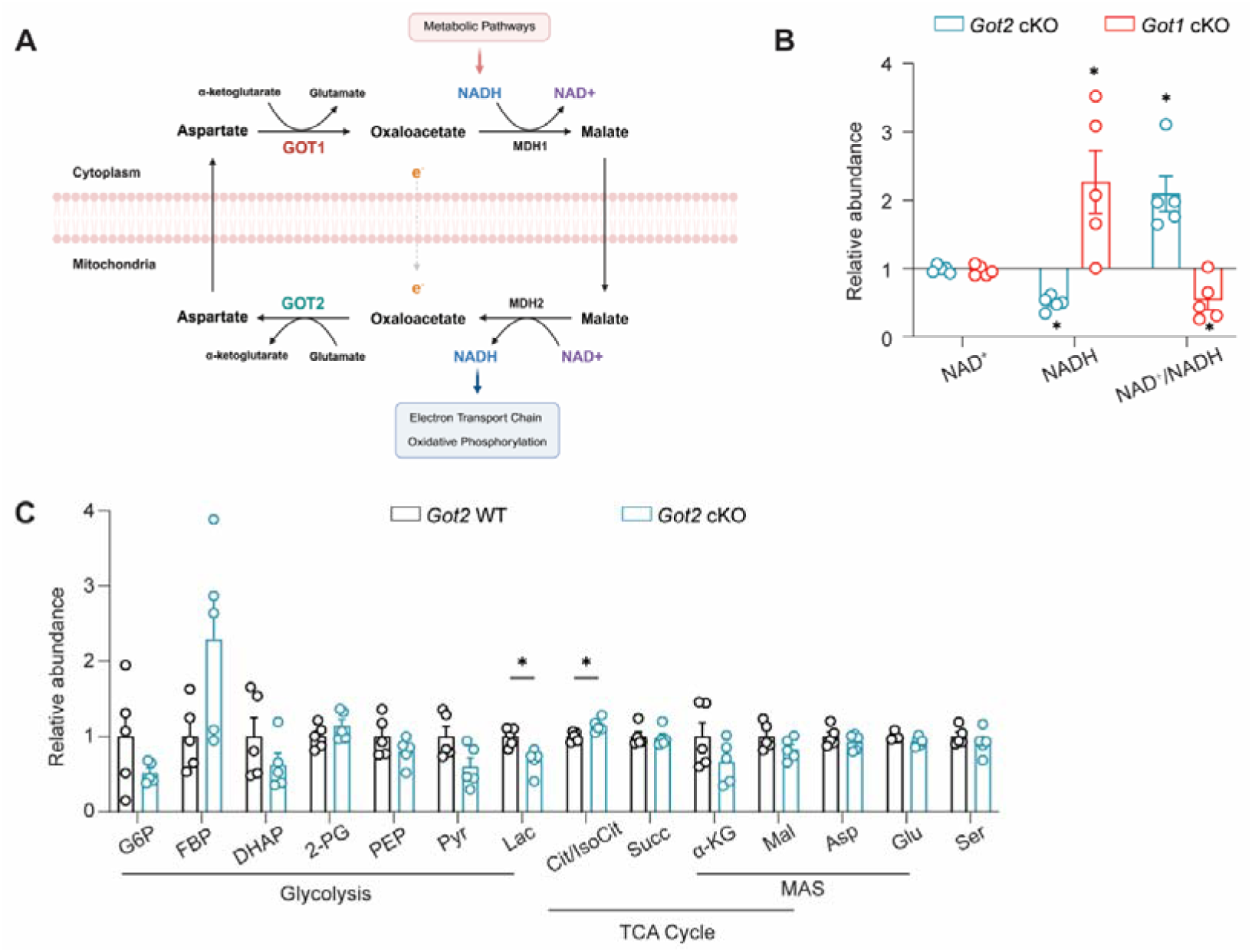
*Got2 cKO* perturbs retinal redox balance differently than *Got1* cKO. **(A)** Schematic showing GOT1 and GOT2 within the malate-aspartate shuttle (MAS) (created with Biorender); **(B)** Relative abundance of NAD^+^ and NADH and the NAD^+^/NADH ratio in the *Got2* cKO and *Got1* cKO retina at 2 months of age (each normalized to respective WT retina); **(C)** Relative abundance of metabolites in glycolysis, the TCA cycle and the MAS in the retina of WT and *Got2* cKO mice. Relative abundance (fold change) was calculated by normalizing ion intensity to the mean value of WT samples. N=5 animals per group; Statistical differences are based on an unpaired two-tail student’s t-test as compared to WT mice; *P <0.05. Graph presents mean ± SEM. NAD^+^: nicotinamide adenine dinucleotide, NADH: nicotinamide adenine dinucleotide + hydrogen, G6P: glucose-6-phosphate, FBP: fructose 1,6-bisphosphatase, DHAP: dihydroxyacetone phosphate, 2-PG: 2-phosphoglycolate, PEP: phosphoenolpyruvate, Pyr: pyruvate, Lac: lactate, Cit: citrate, IsoCit: Isocitrate, Succ: succinate, α-KG: alpha-ketoglutarate, Mal: malate, Asp: aspartate, Glu: glutamate, Ser: serine. Relative abundance data for *Got1* cKO mice was reused with permission. Copyright © 2023 Subramanya, Goswami, Miller, Weh, Chaudhury, Zhang, Andren, Hager, Weh, Lyssiotis, Besirli and Wubben.

### *Got2* cKO induces changes in the expression of metabolism-related genes distinct from those observed in *Got1* cKO retina

The lack of changes in steady-state metabolite levels may be due to rod PRs rewiring their metabolism to maintain homeostasis in the face of GOT1 or GOT2 knockout. To assess this possibility, the expression of genes involved in the MAS, glycolysis, pyruvate metabolism, TCA cycle, amino acid metabolism and redox balance was assessed in 2-month-old *Got2* cKO and WT retina using qRT-PCR (Fig. 3 and Supp. Table. 1). In accordance with GOT2 protein expression (Fig. 1B), *Got2* expression is decreased in the *Got2* cKO retina compared to WT (Fig. 3A) with *Got1* expression showing a small but statistically significant increase. Additionally, the expression of the NAD-dependent malate dehydrogenases (*Mdh1* and *Mdh2*), which are critical components of the MAS, are increased in the *Got2* cKO retina (Fig. 3A). Beyond those crucial to the MAS, the expression of numerous other genes involved in glycolysis (*Pfkl*), pyruvate metabolism (*Pdk3, Pdhb, Me2*), the TCA cycle (*Aco1, Idh3g, Sucla2, Suclg1, Sdha, Sdhb, Sdhc, Sdhd, Fh1*), amino acid metabolism (*Bcat2, Gls2, Asns*), and redox balance (*Gpx1, Gpx4, Pgd, Sod1, Sod2, Cat*) were altered in the 2-month-old *Got2* cKO retina compared to WT (Figs. 3B-F). Certainly, alterations in gene expression do not always equate to protein level changes nor dictate metabolic flux, but the significant changes in retinal metabolism-related gene expression observed here does suggest that retinal metabolic reprogramming upon loss of GOT2 in rod PRs may impact the retinal metabolite changes, or lack thereof, noted at 2 months of age (Fig. 2C). Interestingly, many of the genes that are upregulated upon loss of GOT2 in rod PRs were previously shown to be downregulated in the *Got1* cKO retina compared to WT at 2 months of age (Fig. 3) further illustrating the differential dependence of rod PR metabolism, function and survival on GOT1 versus GOT2 [24].

**Fig. 3.**
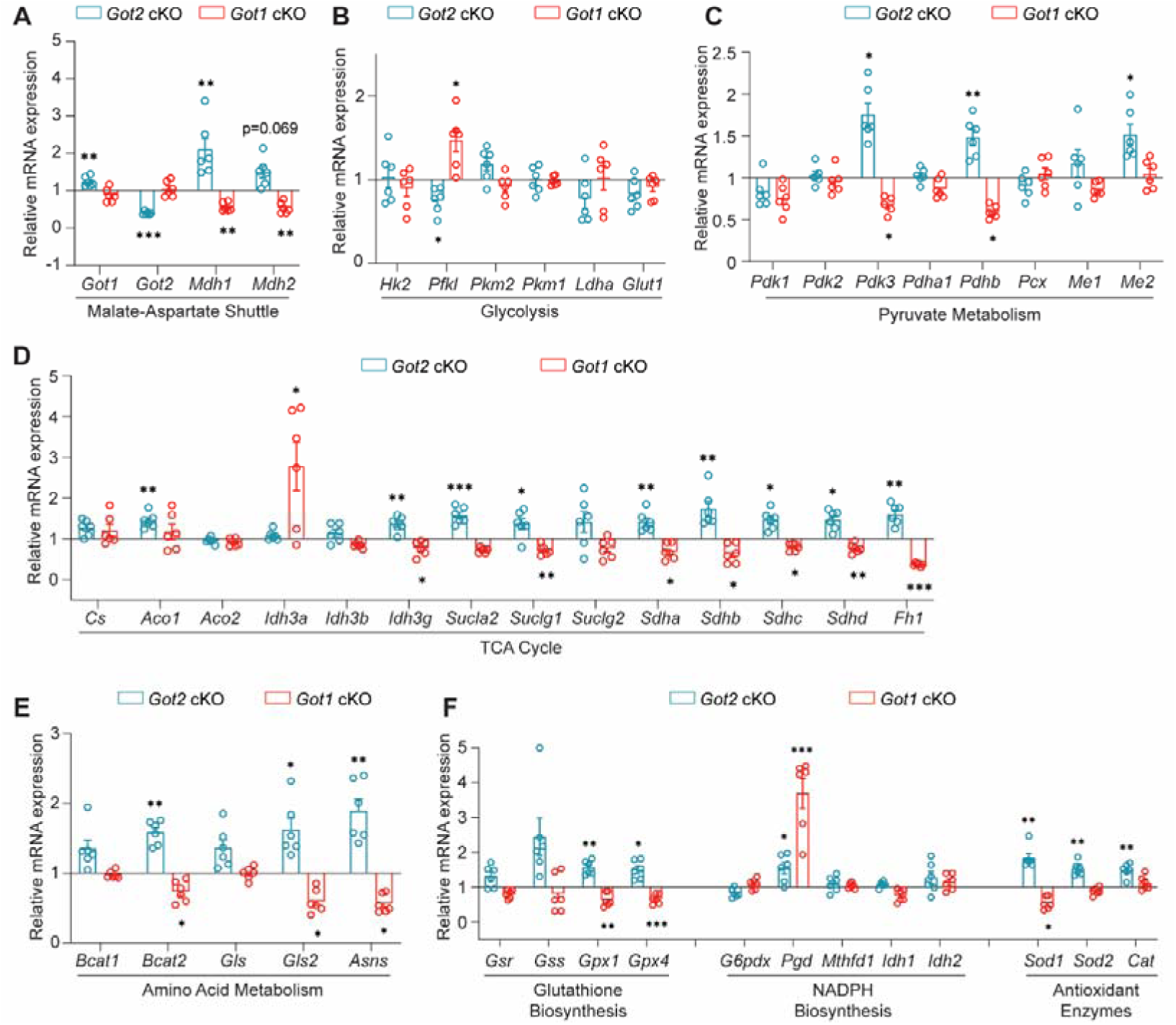
Loss of *Got2* in rod PRs impacts the expression of retinal metabolism-related genes opposite that of *Got1*. qRT-PCR analysis of genes related to **(A)** MAS, **(B)** glycolysis, **(C)** pyruvate metabolism, **(D)** TCA cycle, **(E)** amino acid metabolism, and **(F)** redox balance in 2-month-old *Got2* cKO and *Got1* cKO retina. Relative mRNA expression (fold change) was calculated using the 2^-ΔΔCt^ method and normalized to WT retinas. N=5-6 animals per group. Statistical differences are based on an unpaired two-tail student’s t-test as compared to WT mice; *P <0.05, **P<0.01, ***P<0.001. Mean ± SEM. Relative mRNA expression data for *Got1* cKO mice was reused with permission. Copyright © 2023 Subramanya, Goswami, Miller, Weh, Chaudhury, Zhang, Andren, Hager, Weh, Lyssiotis, Besirli and Wubben.

### *Got2* cKO, unlike *Got1* cKO, does not significantly impact retinal mitochondrial function

The canonical biochemical role of the MAS and its key enzymes, GOT1 and GOT2, is to shuttle reducing equivalents (i.e. NADH) from the cytosol to the mitochondria to support oxidative phosphorylation (Fig. 2A). In line with this, the NAD^+^/NADH ratio was altered in the *Got1* and *Got2* cKO retina as compared to WT, respectively, albeit in opposing directions (Fig. 2B). Thus, we sought to assess how the loss of GOT1 or GOT2 in rod PRs impacts retinal mitochondrial function.

Mitochondrial stress tests were performed on 2-month-old retinas from *Got2* cKO, *Got1* cKO and WT mice using the BaroFuse [50]. The fractional change in oxygen consumption rate (OCR) in response to oligomycin or carbonyl cyanide 4-(trifluoromethoxy)phenylhydrazone (FCCP) was unchanged between the *Got2* cKO and WT 2-month-old retina (Fig. 4A and B). Similarly, the expression of oxidative phosphorylation complexes was unchanged in the *Got2* cKO and WT retina (Fig. 4C). In contrast, *Got1* cKO resulted in a significant decrease in the fractional change in retinal OCR following FCCP treatment as compared to WT (Fig. 4D and E). This observed reduction in maximal respiratory response was not due to alterations in the expression of oxidative phosphorylation components between the *Got1* cKO and WT retina (Fig. 4F). This finding is in accordance with a previous *in vitro* study where GOT1 inhibition and the resultant decrease in cellular NAD^+^/NADH ratio impaired mitochondrial function and further highlights the differential impact of GOT1 and GOT2 knockout on PR metabolism and health [33].

**Fig. 4.**
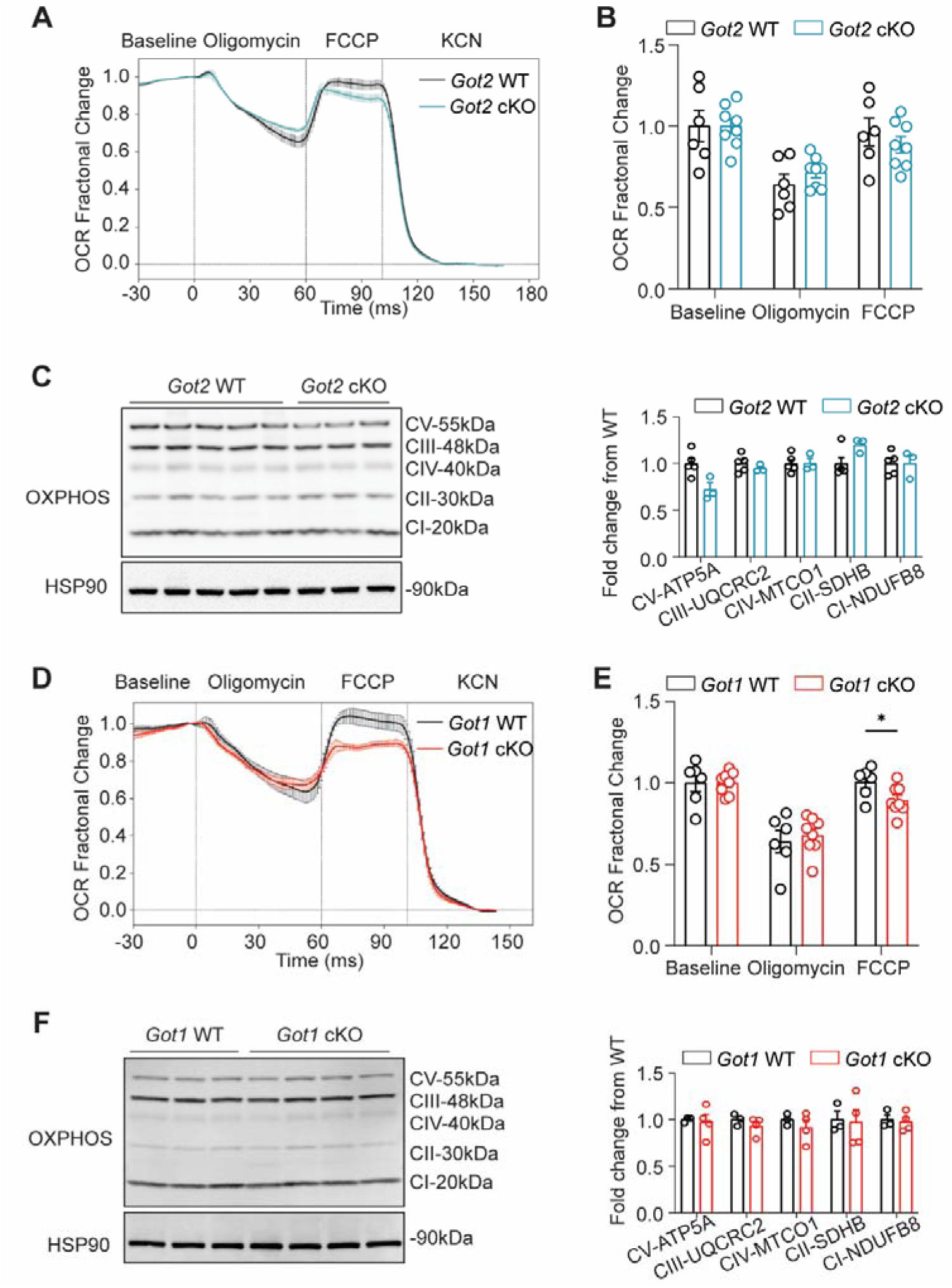
*Got2* cKO, unlike *Got1* cKO, does not significantly impact mitochondrial function. **(A)** Mitochondrial stress test performed on 2-month-old *Got2* cKO and WT retina using the Barofuse. The baseline was established by perifusing the tissue for 90 min prior to oligomycin, FCCP, and KCN being sequentially injected into the perifusate as noted. Data are plotted as fractional change relative to baseline (Mean ± SE). **(B)** Quantitation of the fractional change in oxygen consumption rate (OCR) in WT and *Got2* cKO retina upon the addition of oligomycin and FCCP. N=6-8 animals per group. **(C)** Western blot analysis and quantitation of the mitochondrial oxidative phosphorylation complexes show no differences between 2-month-old WT and *Got2* cKO retina. N=3-5 animals per group. **(D-F)** Same analyses with similar number of animals as in A-C using 2-month-old *Got1* cKO and WT retina. Statistical differences are based on an unpaired two-tail student’s t-test as compared to WT mice; *P <0.05. Mean ± SEM. FCCP: carbonyl cyanide p-trifluoromethoxyphenylhydrazone, KCN: potassium cyanide, CI-NDUFB8: complex 1, NADH:ubiquinone oxidoreductase subunit B8, CII-SDHB: complex 2, succinate dehydrogenase complex iron sulfur subunit B, CIII-UQCRC2: complex 3, ubiquinol-cytochrome c reductase core protein 2, CV-ATP5A: complex 5, ATP synthase F1 subunit alpha, HSP90: heat shock protein 90.

### Redox imbalance is critical to rod PR degeneration following *Got1* cKO

While the conventional paradigm in redox biology emphasizes oxidative stress as a key driver of pathology in numerous diseases including retinal degenerative diseases [53,54], recent evidence indicates that its converse, NADH reductive stress, which is marked by NADH accumulation, is just as detrimental to cell function and survival [55,56]. So, we hypothesized that NADH accumulation contributes to the robust PR degeneration observed in the *Got1* cKO retina. NADH accumulation and the resultant reductive stress can be relieved with electron acceptors, such as pyruvate [32]. Pyruvate accepts electrons from NADH to regenerate NAD^+^ and produces lactate via the enzyme lactate dehydrogenase (Fig. 5A). Therefore, pyruvate supplementation or its increased production would be expected to prolong PR survival in the *Got1* cKO mouse. *Got1* cKO animals were supplemented with 500 mg/kg/d of sodium pyruvate in the drinking water starting at 2 months of age [41] and the thickness of the IS/OS layer, ONL and total retina was monitored over time using OCT. Indeed, systemic pyruvate supplementation produced a significant increase in the ONL and total retinal thickness after 2 months of treatment (4 months of age), restoring 65% of ONL and 40% of total retinal thickness, and after 4 months of treatment (6 months of age), pyruvate supplementation continued to preserve ONL and total retinal thickness with a protective effect observed on the IS/OS as well (Fig. 5B).

**Fig. 5.**
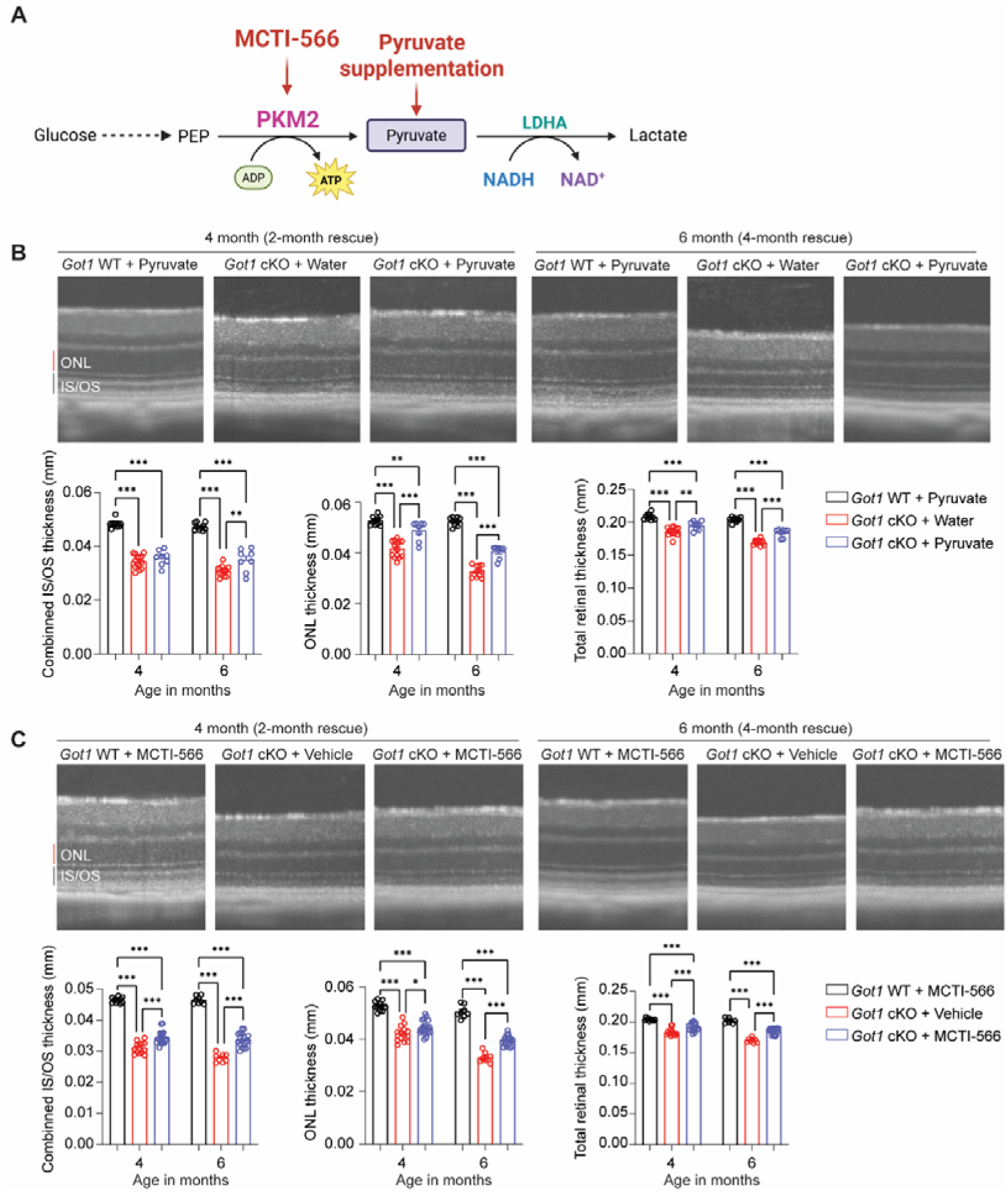
Redox imbalance impacts rod PR degeneration following *Got1* cKO. **(A)** Schematic illustration of redox modulation strategies (created with Biorender). Pyruvate supplementation (oral supplementation from 2-6 months) and a small molecule PKM2 activator, MCTI-566 (i.p., 25 mg/kg biweekly from 2-6 months) was used to theoretically shift the NAD^+^/NADH ratio by promoting pyruvate-to-lactate conversion in *Got1* cKO animals; **(B)** Representative OCT images and related quantitative analysis demonstrate the thickness of IS/OS, ONL, and TRT after 2-month (4 months of age) and 4-month (6 months of age) pyruvate supplementation; N=4-7 animals per group; **(C)** Representative OCT images and related quantitative analysis reveal significant improvement in the thickness of IS/OS, ONL, and TRT after 2-month treatment (4 months of age) and 4-month treatment (6 months of age) with MCTI-566; N=4-9 animals per group. Statistical differences are based on two-way ANOVA; *P <0.05, **P<0.01 and ***P<0.001. Graph presents mean ± SEM. PKM2: pyruvate kinase muscle isoform 2, PEP: phosphoenolpyruvate, NAD^+^: nicotinamide adenine dinucleotide, NADH: nicotinamide adenine dinucleotide + hydrogen, LDHA: lactate dehydrogenase A, IS/OS: inner segment/outer segment, ONL: outer nuclear layer, TRT: Total retinal thickness.

Pyruvate kinase muscle isoform 2 (PKM2) catalyzes the penultimate step in glycolysis, converting phosphoenolpyruvate (PEP) and adenosine diphosphate (ADP) to pyruvate and ATP, with PKM2 predominantly expressed in PRs in the retina. The activity of PKM2 is tightly regulated and governed by its oligomeric state [11]. We recently developed novel, small molecule PKM2 activators, including MCTI-566 [40]. MCTI-566 crosses the blood-retina barrier after systemic administration to activate PKM2 in the retina, is specific for PRs and PKM2 in the retina and does not demonstrate anatomic or functional safety signals. Importantly, activating PKM2 is PR neuroprotective in preclinical models of PR degeneration [17,40,57,58]. As MCTI-566 is highly selective for PKM2, with PKM2 predominantly expressed in PRs, MCTI-566 may provide PR specificity for increasing pyruvate production. By enhancing pyruvate production and its subsequent conversion to lactate, PKM2 activation by MCTI-566 may facilitate NADH oxidation, thereby alleviating reductive stress in PRs (Fig. 5A). Based on our previously published pharmacokinetic analyses [40], *Got1* cKO mice were injected intraperitoneally (IP) every other week with MCTI-566 (25 mg/kg) starting at 2 months of age, which is a time prior to PR loss. MCTI-566 treatment resulted in preservation of total retinal (40%, 2 months; 47%, 4 months), ONL (22%, 2 months; 37%, 4 months), and IS/OS (21%, 2 months; 31%, 4 months) thickness compared to vehicle-treated *Got1* cKO mice with the neuroprotective effect of PKM2 activation increasing with treatment duration (Fig. 5C).

The rescue of ONL and IS/OS thickness with pyruvate supplementation and MCTI-566 treatment, coupled with the fact that NAD^+^ level was unchanged in the *Got1* retina compared to WT (Fig. 2B), suggests the neuroprotective mechanism is dependent on NADH turnover and not NAD^+^ synthesis. To test this further, we expressed cytosolic NADH oxidase from *Lactobacillus brevis* (*Lb*NOX) in rod PRs of the *Got1* cKO mouse. *Lb*NOX oxidizes NADH to NAD^+^ using molecular oxygen (Fig. 6A). The previously published *cytoLbNOX^LSL/LSL^* mouse was crossed to the *Got1* cKO (*cytoLbNOX^LSL/LSL^ ;Got1^fl/fl^*;*Rho-Cre^+^; cytoLbNox/Got1* cKO) and *Got1* WT (*cytoLbNOX^LSL/LSL^ ;Got1^wt/wt^*;*Rho-Cre^+^; cytoLbNox/Got1* WT) mouse, both of which express a Cre-recombinase under the control of the rhodopsin promoter [39]. In the *cytoLbNOX^LSL/LSL^*mouse, a Lox-P-flanked STOP codon is upstream of *cytoLbNOX,* which is followed by an internal ribosome entry site (IRES)-linked enhanced green fluorescent protein (eGFP) sequence (Fig. 6B). Therefore, in the presence of a Cre-recombinase, the upstream STOP codon is excised and cytosolic *Lb*NOX is expressed along with eGFP. Western blot analysis demonstrates eGFP protein expression, a surrogate for *Lb*NOX expression, in total retinal lysate from *cytoLbNox/Got1* WT and *cytoLbNox/Got1* cKO mice as compared to *Got1* WT mice without cytosolic *Lb*NOX (Fig. 6C). Furthermore, representative immunofluorescence images show eGFP expression is restricted to the PR layer in the retina (Fig. 6D). Expression of cytosolic *Lb*NOX in *Got1* WT retina did not impact the IS/OS, ONL or total retinal thickness as compared to WT mice that did not express the cytosolic *Lb*NOX (Fig. 6E). Importantly, in line with our hypothesis, cytosolic *Lb*NOX expression in *Got1* cKO rod PRs improved IS/OS thickness as compared to *Got1* cKO mice that do not express the cytosolic *Lb*NOX and restored 73% of ONL thickness at 4 months of age (Fig. 6F). These findings provide strong evidence via multiple orthogonal approaches that NADH accumulation and reductive stress is critical to the PR degeneration observed in the *Got1* cKO mouse.

**Fig. 6.**
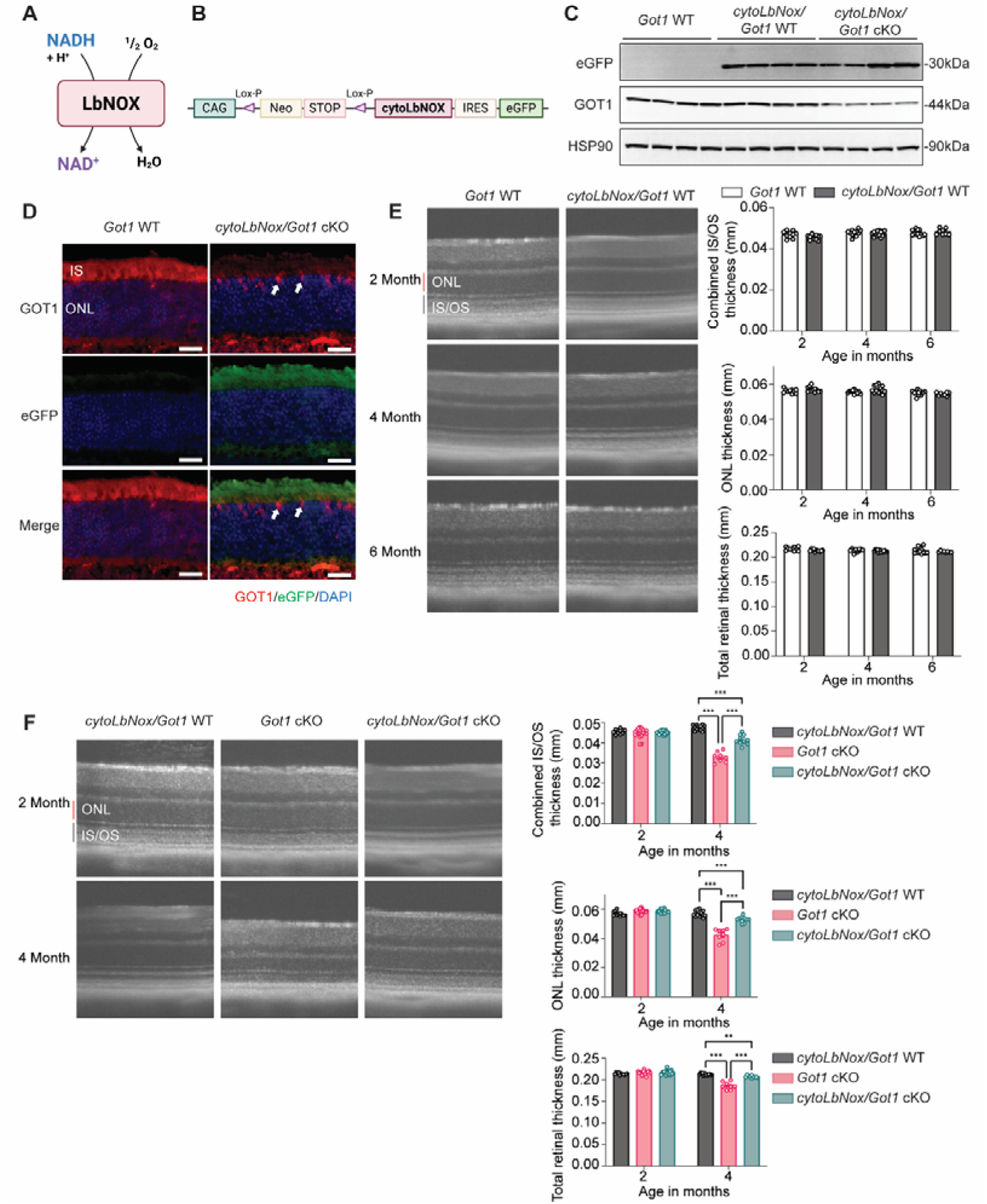
Expression of cytosolic *Lb*NOX in rod PRs rescues PR degeneration in the *Got1* cKO mouse. **(A)** Schematic showing the reaction catalyzed by *Lb*NOX; **(B)** Schematic illustrating the gene targeting strategy used to produce the *cytoLbNOX^LSL/LSL^* mouse strain; **(C)** Western blot analysis confirms the expression of eGFP in total retinal lysate from *cytoLbNox/Got1* WT and *cytoLbNox/Got1* cKO mice as compared to *Got1* WT mice without cytosolic *Lb*NOX, and decreased GOT1 expression level in *cytoLbNox/Got1* cKO retinas; **(D)** Representative immunofluorescence images showing eGFP (green) expression, which is a proxy for the expression of cytosolic *Lb*NOX, and GOT1 (red) expression, which is restricted to cone PRs (white arrow) in *cytoLbNox/Got1* cKO mice; Scale bar: 25 µm; **(E)** Representative OCT images and related quantitative analysis reveal no significant differences in IS/OS, ONL, or total retinal thickness at 2, 4, and 6 months of age between *cytoLbNox/Got1* WT and *Got1* WT mice without cytosolic *Lb*NOX; N=4-7 animals per group. **(F)** Representative OCT images and related quantitative analyses show significantly improved IS/OS, ONL, and total retinal thickness in *cytoLbNox/Got1* cKO mice as compared to *Got1* cKO mice without cytosolic *Lb*NOX at 4 months of age. N=4-9 animals per group. Statistical differences in (E) are based on an unpaired two-tail student’s t-test and (F) are based on two-way ANOVA; **P<0.01 and ***P<0.001. Graph presents mean ± SEM. NAD^+^: nicotinamide adenine dinucleotide, NADH: nicotinamide adenine dinucleotide + hydrogen, IRES: internal ribosome entry site, eGFP: enhanced green fluorescent protein, IS/OS: inner segment/outer segment, ONL: outer nuclear layer. Schematics created with Biorender.

### GOT1 and GOT2 are regulated differently during PR stress

A previous study utilizing a model of autosomal recessive RP suggested that the GOTs may be biomarkers of early PR degeneration and potential PR neuroprotective targets [35]. To provide greater insight into the relevance of these enzymes in retinal degenerative diseases, we assessed GOT1 and GOT2 protein expression in preclinical models of inherited and acquired PR stress. Retinas were harvested from *rd10*, *rd2* and *rd12* mice at ages prior to the onset of PR degeneration. These preclinical models of inherited PR degeneration result from perturbations in three different genes and represent early-onset rod degeneration (*rd10*, *Pde6b*) [59], rod-cone degeneration (*rd2*, *Prph2*) [60] and slow PR degeneration secondary to retinal pigment epithelium (RPE) dysfunction (*rd12, Rpe65*) [61]. Interestingly, GOT2 protein expression was decreased in the retina from all three models of inherited PR degeneration as compared to WT but GOT1 levels were unchanged in comparison to WT (Fig. 7A-C). To assess if these alterations in GOT1 and GOT2 are specific to inherited PR degeneration models or also occur in models of acquired PR stress, experimental retinal detachment was performed in WT mice and retinas harvested 1, 3 and 7 days after detachment. Retinal detachment (RD) separates the PRs from the underlying RPE and choroid, so the supply of oxygen, glucose, and other nutrients to the PRs is disrupted leading to PR degeneration over time [62,63]. GOT1 protein expression did not change significantly over time while GOT2 expression was decreased starting at 3 days post detachment (Fig. 7D), a time prior to significant PR degeneration [64]. Of note, the expression of complexes involved in oxidative phosphorylation including Complex V (ATP5A), Complex III (UQCRC2), Complex IV (MTCO1), Complex II (SDHB), and Complex I (NDUFB8), as well as TIM23, a protein anchored in the inner mitochondrial membrane, were unchanged in these models of PR stress. These data suggest the down regulation of the mitochondrially-located GOT2 is not attributable to a general reduction in mitochondrial content (Supp. Fig. 2).

**Fig. 7.**
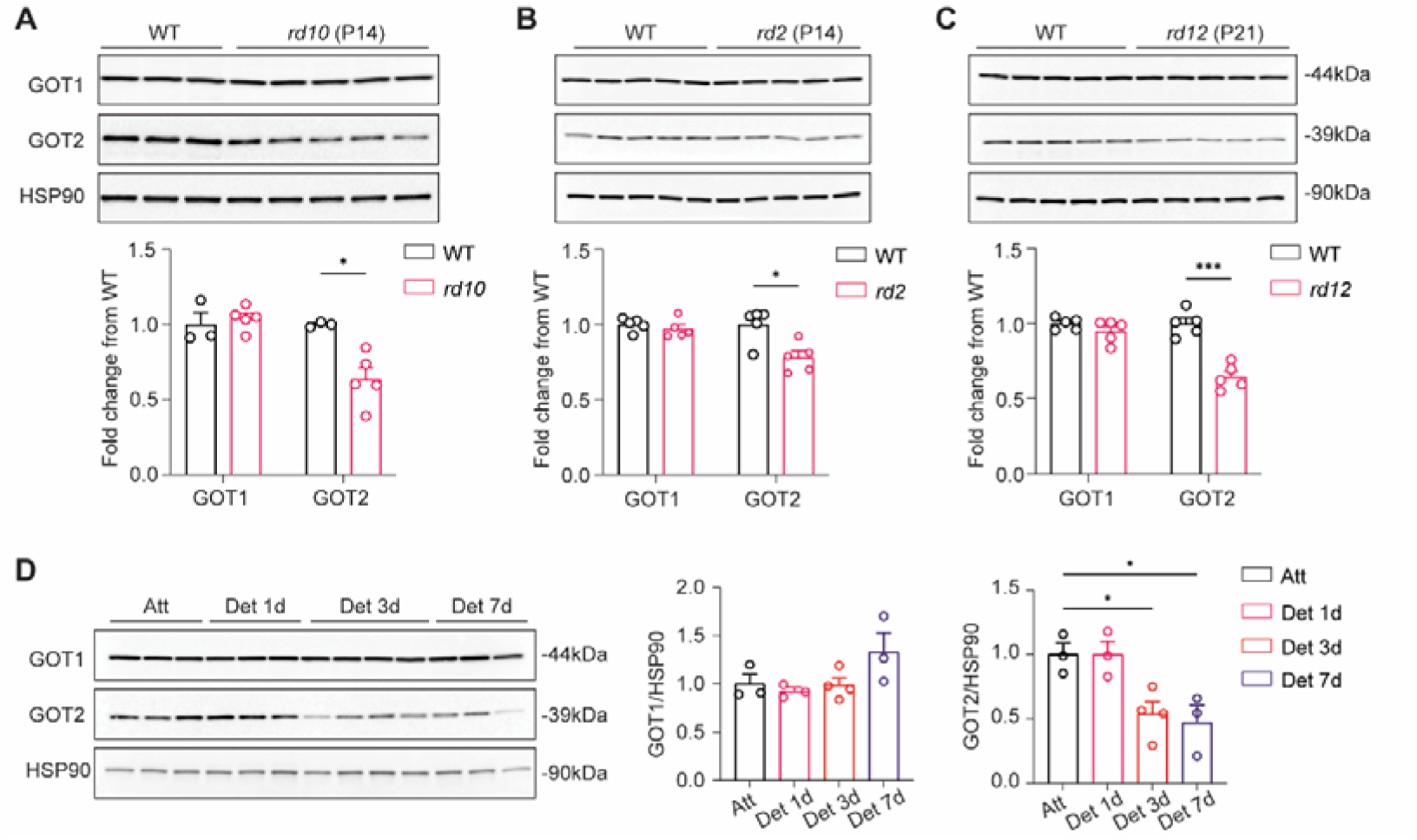
Retinal GOT1 and GOT2 are regulated differently in preclinical models of PR stress. **(A-C)** Western blot analysis and quantitation of GOT1 and GOT2 from *rd10* (P14), *rd2* (P14), and *rd12* (P21) retina as compared to age-matched wild-type (WT) retina. **(D)** Western blot analysis and quantitation of GOT1 and GOT2 in attached (Att) as well as 1-day, 3-day, and 7-day detached (Det) retina. N=3-5 animals per group; Statistical differences in (A-C) are based on an unpaired two-tail student’s t-test, and (D) are based on one-way ANOVA as compared to WT retina; *P <0.05 and ***P<0.001. Mean ± SEM. HSP90: heat shock protein 90.

### Loss of *Got2* promotes PR resistance to acute stress in the experimental retinal detachment model

The GOTs are important mediators of redox balance through their impact on the NAD^+^/NADH ratio (Fig. 2A and B). As GOT2, but not GOT1, is decreased in multiple models of PR degeneration (Fig. 7), we next sought to determine if an altered retinal NAD^+^/NADH ratio is an early feature in one of these models, experimental retinal detachment. Indeed, 24 h after experimental retinal detachment in rats or mice, the retinal NAD^+^/NADH ratio was statistically significantly decreased in detached as compared to attached rat retinas with a downward trend observed in detached mouse retinas as well (Fig. 8A). Notably, this trend in mouse retina is likely secondary to NADH accumulation rather than a reduction in the NAD^+^ level (Supp. Fig. 3).

**Fig. 8.**
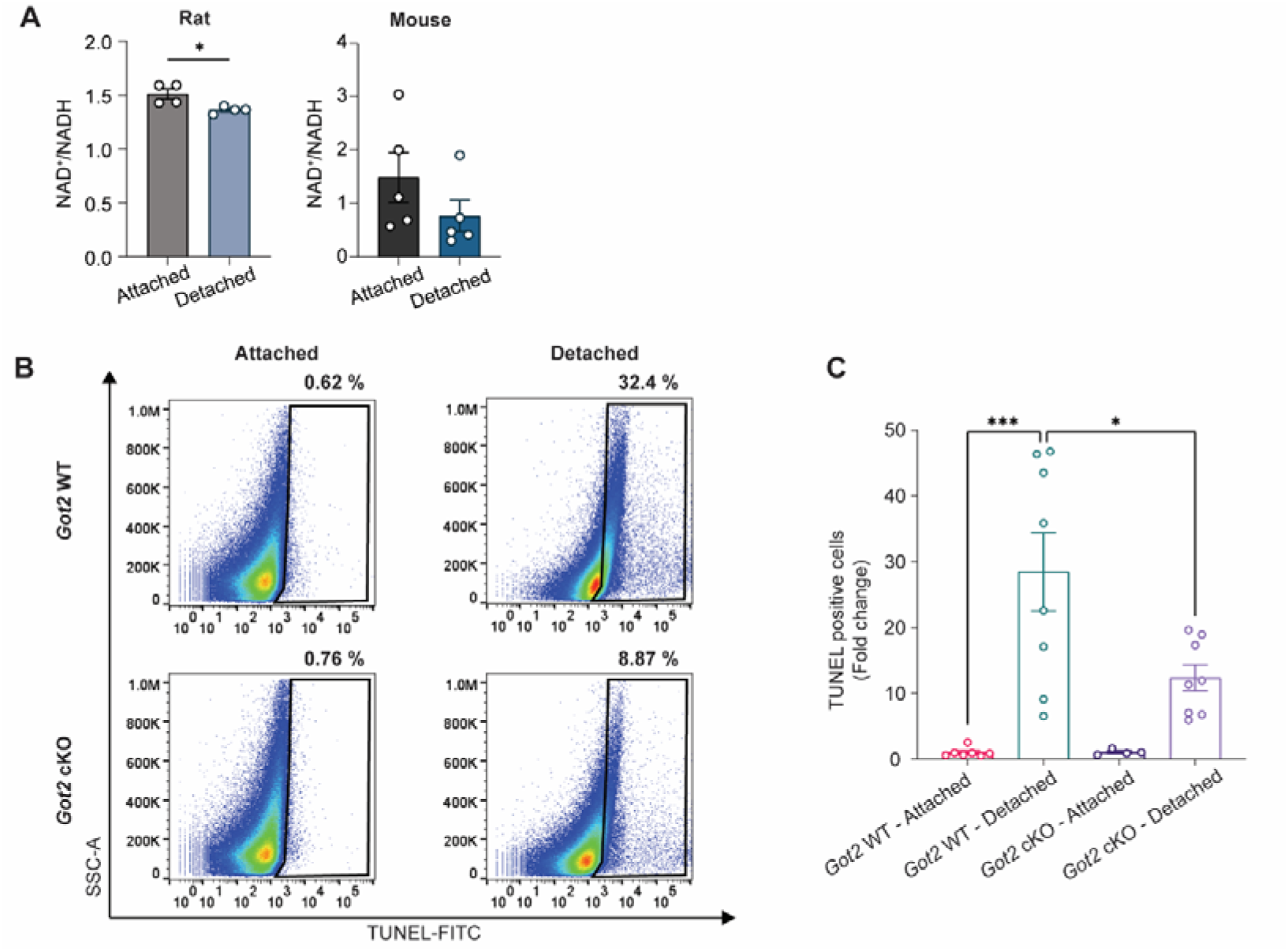
*Got2* cKO demonstrates photoreceptor neuroprotection in retinal detachment model. **(A)** NAD^+^/NADH ratio is decreased in 24-hour detached retina in both rats and mice; N=4-5 rat retinas per group; for mouse, attached or detached retinas were respectively pooled across two different animals for each N with N=5 for both the attached and detached mouse groups; **(B)** Representative flow cytometry contour plots to assess TUNEL-positive cells in WT (top) or *Got2* cKO (bottom) retinas three days after experimental retinal detachment. The black gate represents TUNEL-positive cells; **(C)** Fold change in the percent of TUNEL-positive cells in the retina normalized to *Got2* WT attached retina. N=4-9 retinas per group. Statistical differences in (A) are based on an unpaired two-tail student’s T-test and (C) are based on one-way ANOVA; *P <0.05 and ***P<0.001. NAD^+^: nicotinamide adenine dinucleotide, NADH: nicotinamide adenine dinucleotide + hydrogen.

The experimental retinal detachment model of PR degeneration has been used extensively to identify factors and assess therapeutic strategies that promote PR resistance to stress [17,42,63,65,66]. Considering GOT2 expression is decreased after retinal detachment (Fig. 7D) and GOT2 knockdown in rod PRs impacts the retinal NAD^+^/NADH ratio in a direction opposite to reductive stress and that observed after retinal detachment (Fig. 2B and 8A), we proceeded to assess the effect of *Got2* cKO on PR apoptosis during retinal detachment. Retinal detachment was induced in 2-month-old *Got2* cKO and WT animals and whole retina was collected 3 days after detachment for TUNEL analysis via flow cytometry. PR TUNEL staining peaks at this time point and correlates with long-term PR survival [64]. Retinal detachment in *Got2* cKO mice resulted in more than a 50% decrease in TUNEL-positive cells compared to *Got2* WT mice (Fig. 8B and C). Hence, *Got2* cKO demonstrates PR neuroprotection in this model of acute outer retinal stress.

## Discussion

While GOT1 and GOT2 are both essential MAS components, in this study we show a differential dependence on the respective isoforms for rod PR health at baseline and during stress. Under non-stress *in vivo* conditions, *Got1* cKO causes PR degeneration and is accompanied by NADH accumulation and a decreased retinal NAD^+^/NADH ratio [24]. Here, we show that NADH oxidation via metabolic or genetic means prolongs PR survival in *Got1* cKO animals, implicating NADH accumulation, or reductive stress, as a key driver of PR degeneration. In contrast to the *Got1* cKO, *Got2* cKO causes minimal PR degeneration and alterations in retinal NADH and the NAD^+^/NADH ratio that oppose reductive stress at baseline. Interestingly, GOT2, but not GOT1, is decreased in multiple models of PR degeneration, including RD where the NAD^+^/NADH ratio favors a reductive state. Notably, the loss of GOT2 in PRs demonstrates a neuroprotective effect after experimental RD suggesting decreased GOT2 expression may be part of a PR stress response and a novel neuroprotective target that combats NADH reductive stress to prolong PR survival in retinal degenerative disease.

The biosynthetic demand of PRs rivals that of cancer cells despite being terminally differentiated [67], and similarities exist between cancer cell and PR metabolism [12,67,68]. *In vitro* studies in pancreatic cancer cells have shown that disrupting either *Got1* or *Got2* impairs cell proliferation and colony formation [31–33]. Accordingly, we previously demonstrated loss of *Got1* in rod PRs *in vivo* results in robust PR degeneration [24]. Considering these *in vivo* results, the aforementioned *in vitro* studies where disrupting either *Got1* or *Got2* results in a similar phenotype, and that GOT1 and GOT2 are both critical components of the MAS, the rather modest retinal degenerative phenotype observed with conditional deletion of *Got2* in rod PRs *in vivo* was unexpected (Fig. 1). A recent report has demonstrated, that in contrast to *in vitro* studies, the GOTs are dispensable for pancreatic cancer development *in vivo* [69]. As such, our data adds to a growing body of literature that emphasizes the importance of the *in vivo* context when studying these metabolic enzymes [32,69] and further demonstrates the role of tissue/cell type in dictating the dependence on the GOTs [51].

The GOTs within the MAS are important mediators of redox balance and cellular metabolism through their impact on the NAD^+^/NADH ratio (Fig. 2A). Accordingly, loss of either *Got1* or *Got2* in rod PRs altered the retinal NAD^+^/NADH ratio, albeit in opposite directions (Fig. 2B). An elevated retinal NAD^+^/NADH ratio was observed in the *Got2* cKO retina secondary to a decrease in the relative abundance of NADH. In contrast, the *Got1* cKO retina demonstrated an accumulation of NADH and a decreased retinal NAD^+^/NADH ratio. Taking into account this stark difference in the NAD^+^/NADH ratio when *Got1* or *Got2* was conditionally knocked out in rod PRs and the fact that the *Got1* cKO mouse demonstrated much more robust PR degeneration as compared to the *Got2* cKO mouse (Fig. 2) [24], we hypothesized that NADH reductive stress, marked by NADH accumulation, is critical to the PR degeneration observed in the *Got1* cKO mouse. Reductive stress can impact cell survival and function through ER protein misfolding, mitochondrial dysfunction and inhibition of critical metabolic pathways [55,56], all of which have been shown to play a role in PR degeneration [3,8,70,71]. Of note, mitochondrial dysfunction was observed in the *Got1* cKO but not in the *Got2* cKO retina (Fig. 5). Moreover, our hypothesis was supported by data illustrating that supplementation of an electron acceptor like pyruvate, pharmacologic treatment with a PKM2 activator to increase pyruvate production or expression of a cytosolic NADH oxidase prolongs PR survival in the *Got1* cKO mouse (Figs. 5 and 6).

Pyruvate can affect the redox state of the cell via its reduction by lactate dehydrogenase (Fig. 5A) but also can support the TCA cycle or act as a biosynthetic substrate to induce aspartate synthesis [72]. With regards to these latter metabolic processes, TCA cycle metabolites were not significantly altered in the *Got1* cKO retina, and aspartate levels were actually increased in accordance with GOT1 suppression [24]. So, pyruvate’s impact on the TCA cycle and aspartate biosynthesis may not be critical to its ability to rescue *Got1* cKO PR degeneration. Future metabolomic studies that utilize stable isotope tracing methodologies with uniformly labeled ^13^C-pyruvate (supplementation) or ^13^C-glucose (MCTI-566 treatment) can assess how these rescue strategies are altering metabolism to protect *Got1* cKO PRs.

Additionally, previous *in vitro* studies suggest that GOT1 inhibition promotes the death of certain cancer cells by potentiating ferroptotic stimuli, such as inhibition of cystine import, glutathione biosynthesis or glutathione peroxidase 4 (GPX4) [33]. Interestingly, the *Got1* cKO retina demonstrated a decrease in the expression of numerous redox homeostasis genes, including *Gpx4* (Fig. 3E), as well as key ferroptosis suppressor genes, *Gch1* and *Fth1* [24,73,74]. Future rescue studies with antioxidants, such as N-acetylcysteine, will help illuminate the role of oxidative stress and ferroptosis in this mouse model. That said, rod PR-specific expression of cytosolic *Lb*NOX provided significant rescue of PR degeneration at 4 months of age (Fig. 6). This data strongly implicates NADH reductive stress as a major driving force behind the observed PR loss in the *Got1* cKO mouse as *Lb*NOX directly targets NADH.

Alterations in the NAD^+^/NADH ratio, and NADH reductive stress in particular, have been shown to drive cellular metabolic reprogramming by perturbing the flux of metabolic reactions [56]. Yet, as noted above, few changes in the relative abundance of glycolytic and TCA cycle metabolites were observed (Fig. 2C) [24] despite significant changes in the NAD^+^/NADH ratio when *Got1* or *Got2* was conditionally knocked out in rod PRs. In the transgenic mouse models utilized herein, PRs have been shown to genetically rewire their metabolism in an attempt to maintain metabolic homeostasis and stave off degeneration [11,24,46]. To this end, both the *Got1* cKO and *Got2* cKO retina demonstrated significant changes in the expression of genes involved in the MAS, glycolysis, pyruvate metabolism, the TCA cycle, and amino acid metabolism as compared to WT retina. Moreover, many of the metabolic genes upregulated in the *Got2* cKO retina were downregulated in the *Got1* cKO retina (Fig. 3). These distinct alterations in gene expression may dictate flux through different retinal metabolic pathways to maintain relative metabolite levels and potentially impact the observed retinal NAD^+^/NADH ratio in the *Got1* and *Got2* cKO retina, respectively. Additionally, this complex intracellular rewiring in the context of the outer retinal microenvironment has the potential to impact the observed *in vivo* differences in redox imbalance and retinal degenerative phenotype between the *Got1* and *Got2* cKO mouse models. For example, GOT2 is dispensable in pancreatic tumor formation *in vivo* as cancer-associated fibroblasts secrete pyruvate, providing an electron acceptor to GOT2 KO cancer cells. In turn, pancreatic cancer cells have upregulated pyruvate import and increased macropinocytosis to improve aspartate acquisition and survival [32,75]. Thus, similar metabolic rewiring strategies in PRs *in vivo* could result in their ability to compensate for an impaired MAS, especially in the context of the *Got2* cKO mouse, which shows minimal retinal degeneration over time (Fig. 1). Nonetheless, the fact that loss of *Got2* in rod PRs impacts the expression of retinal metabolism-related genes opposite the loss of *Got1* further emphasizes the observed differential dependence of rod PR metabolism, function and survival on the GOTs.

The role of oxidative stress in PR degeneration and retinal degenerative diseases has been extensively studied [53,54], and in more recent years, several studies have reported on the role of NAD^+^ biosynthesis in PR degeneration [21,76,77]. However, the impact of NADH reductive stress and its manipulation on PR degeneration is poorly understood. Here, we have begun to address this critical knowledge gap by demonstrating that NADH reductive stress is critical to PR degeneration in the *Got1* cKO mouse (Figs. 5 and 6) and may play a role in the experimental RD model (Fig. 8). In this latter model of acute outer retinal stress and PR degeneration, the retinal NAD^+^/NADH ratio is altered in the same direction as in the *Got1* cKO retina to favor reductive stress (Fig. 8A). Hypoxia and mitochondrial dysfunction can both generate NADH reductive stress [56], and both have been observed in the experimental RD model [66,78]. Moreover, similar to its impact in the *Got1* cKO mouse, activating PKM2, such as with MCTI-566, is PR neuroprotective in the experimental RD model [17,40,57]. Hence, NADH reductive stress may play a role in PR degeneration after RD.

Interestingly, though, it is GOT2, but not GOT1, expression that decreases after retinal detachment (Fig. 7D), and *Got2* cKO provided neuroprotection after experimental RD (Fig. 8B and C). The *Got2* cKO mouse causes alterations in retinal NADH and the NAD^+^/NADH ratio that oppose reductive stress (Fig. 2B). As such, decreased GOT2 expression may be part of a PR stress response that combats NADH reductive stress to prolong PR survival in this preclinical model of PR degeneration and potentially others. To this end, GOT2 was decreased at a time prior to significant PR degeneration in multiple preclinical models of IRD regardless of mutation (Fig. 7A-C), and a previous study noted it to be decreased in a separate model of autosomal recessive RP [35]. While future studies need to establish a causal relationship between NADH reductive stress and PR degeneration in these IRD models, mitochondrial dysfunction that can generate reductive stress has been noted in these same or similar genetic models [70,79,80], and activating PKM2 with small molecules, such as MCTI-566, is PR neuroprotective in the *rd10* mouse and trends towards PR neuroprotection in the *rd2* mouse [40,58].

In summary, this study demonstrates the differential dependence on the aspartate aminotransferases, GOT1 and GOT2, for rod PR metabolism, function and survival with loss of GOT2 in rod PRs demonstrating a more modest retinal phenotype as compared to that of GOT1 [24]. This difference in phenotype is likely secondary to redox imbalance with NADH reductive stress driving PR degeneration in the *Got1* cKO retina. Much of the work to date has studied the role of redox imbalance in PR degeneration and retinal degenerative disease through the lens of oxidative stress [53,54]. This study suggests its converse, NADH reductive stress, is also a critical mediator of PR degeneration and targeting GOT2 in preclinical models of retinal degenerative disease may counter this redox imbalance to promote PR resistance to stress. This study provides a strong foundation on the role of NADH reductive stress in PR degeneration and GOT2 as a potential novel neuroprotective target but also opens up key mechanistic questions. For example, a causal relationship between NADH reductive stress and PR degeneration remains to be established in preclinical models beyond RD, and the downstream effectors of such stress that drive disease progression remain unresolved. Understanding the role of NADH accumulation versus NAD^+^ depletion in retinal degenerative diseases will be critical as well. By improving our understanding of NADH reductive stress and its molecular mechanisms in retinal degenerative disease, we expect to identify novel targets, such as GOT2, with far-reaching therapeutic potential.

## Acknowledgements

Funding for this research was supported by a NEI K08EY031757 and a RPB Unrestricted Grant to the Department of Ophthalmology and Visual Sciences at the University of Michigan, Kellogg Eye Center. This work utilized the Vision Research Core funded by P30 EY007003 from the National Eye Institute and P30 DK020572 from the National Institute of Diabetes and Digestive and Kidney Diseases. CAL was supported by NCI grants R37CA237421, R01CA244931 and R01CA248160. RCR was supported by an OHSU Foundation Grant, an unrestricted grant from Research to Prevent Blindness, New York, NY, NIH P30 EY010572 Core Grant, and the Malcolm M. Marquis, MD Endowed Fund for Innovation to Casey Eye Institute, Oregon Health & Science University.

## Conflicts of Interest

TJW is a consultant for Insmed and has equity interest in Ocutheia, Inc. In the past three years, CAL has consulted for Odyssey Therapeutics and Guidepoint Global, and is an inventor on patents pertaining to Kras regulated metabolic pathways, redox control pathways in pancreatic cancer, and targeting the GOT1-ME1 pathway as a therapeutic approach (US Patent No: 2015126580-A1, 05/07/2015; US Patent No: 20190136238, 05/09/2019; International Patent No: WO2013177426-A2, 04/23/2015).

## Authorship Contribution

Meini Chen: Formal analysis, Investigation, Visualization, Writing-Original draft, Writing-review and editing. Eric Weh: Formal analysis, Supervision, Investigation, Writing-Original draft, Writing-review and editing. Moloy T. Goswami: Formal analysis, Investigation, Writing-Original draft, Writing-review and editing. Katherine M. Weh: Writing-Original draft, Writing-review and editing. Heather Hager: Investigation. Peter Sajjakulnukit: Investigation, Resources. Avi Weingarten: Investigation. Shubha Subramanya: Investigation. Nicholas Miller: Investigation. Sraboni Chaudhury: Investigation. Emma Piraino: Investigation. Navdeep S. Chandel: Resources. Renee C. Ryals: Resources, Writing-review and editing. Costas A. Lyssiotis: Conceptualization, Resources, Supervision, Methodology. Thomas J. Wubben: Conceptualization, Formal analysis, Supervision, Funding acquisition, Visualization, Methodology, Writing-Original draft, Writing-review and editing.

## Supplementary

**Supp. Fig. 1:**
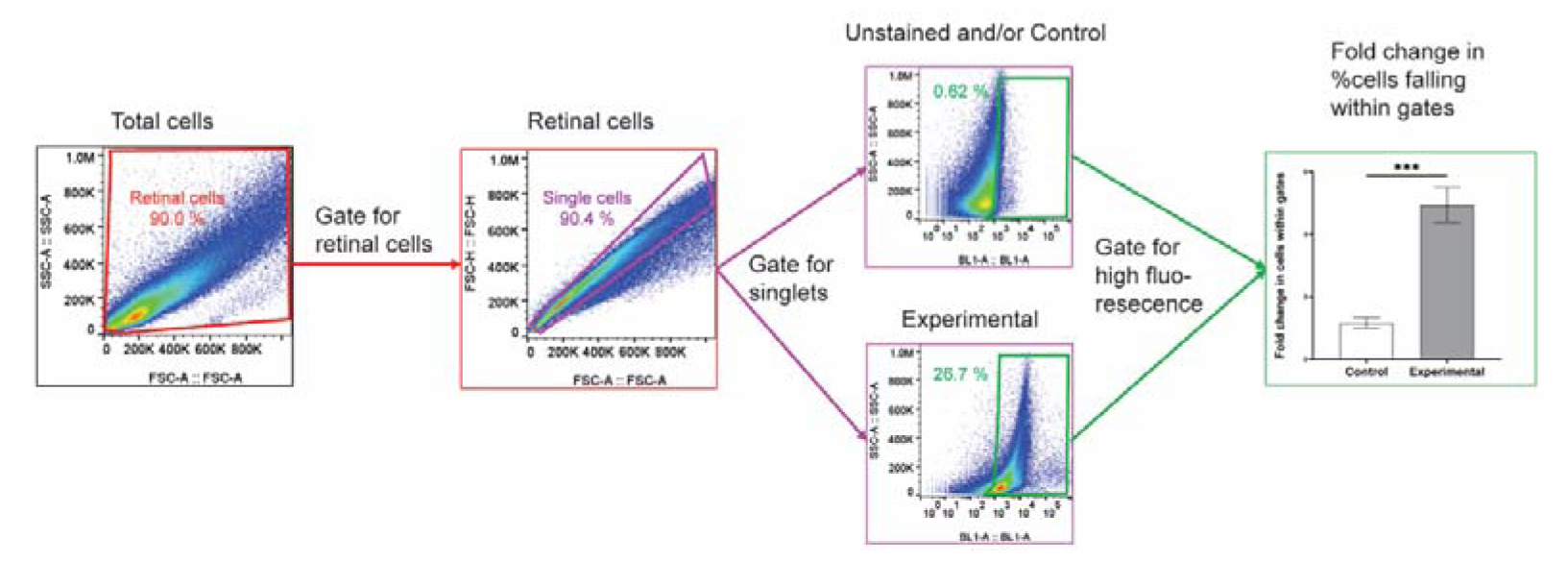
Flow cytometry gating strategy. Cells were first gated by forward and side scatter to exclude debris and broken cells (red gate), followed by selecting singlets using an FSC-A *vs.* FSC-H plot (purple gate). Singlets within the purple gate were then analyzed for a change in fluorescent intensity for each sample (green gate). The fold change in the percentage of cells within the green gate was used to determine changes in TUNEL staining.

**Supp. Fig. 2.**
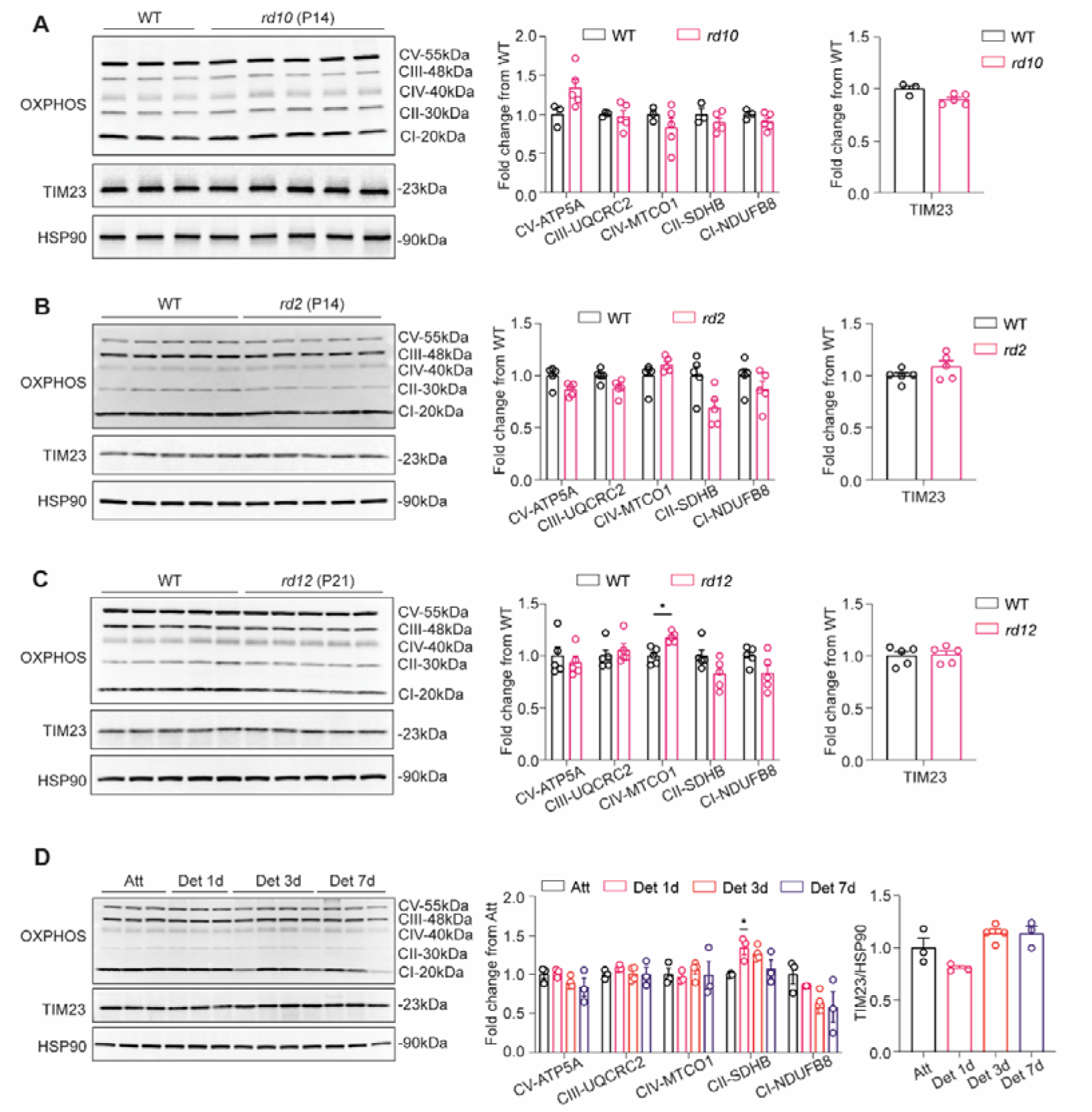
Retinal mitochondrial oxidative phosphorylation complexes and TIM23 remain unchanged in different PR stress models. **(A-C)** Western blot analysis and quantitation of oxidative phosphorylation (OXPHOS) complexes (CV-ATP5A, CIII-UQCRC2, CIV-MTCO1, CII-SDHB, CI-NDUFB8) and TIM23 show no differences between *rd10* (P14), *rd2* (P14), *rd12* (P21) and age-matched wild-type (WT) retina. **(D)** No significant downregulation was observed in the expression of OXPHOS complexes and mitochondrial protein TIM23 in the retinas 1, 3, and 7 days after retinal detachment (Det) compared to attached (Att) retinas. N=3-5 animals per group; Statistical differences in (A-C) are based on an unpaired two-tail student’s T-test, and (D) are based on one-way ANOVA as compared to WT retina; *P <0.05. Mean ± SEM. CI-NDUFB8: complex 1, NADH:ubiquinone oxidoreductase subunit B8; CII-SDHB: complex 2, succinate dehydrogenase complex iron sulfur subunit B; CIII-UQCRC2: complex 3, ubiquinol-cytochrome c reductase core protein 2; CV-ATP5A: complex 5, ATP synthase F1 subunit alpha; TIM23: translocase of the inner membrane 23; HSP90: heat shock protein 90.

**Supp. Fig. 3.**
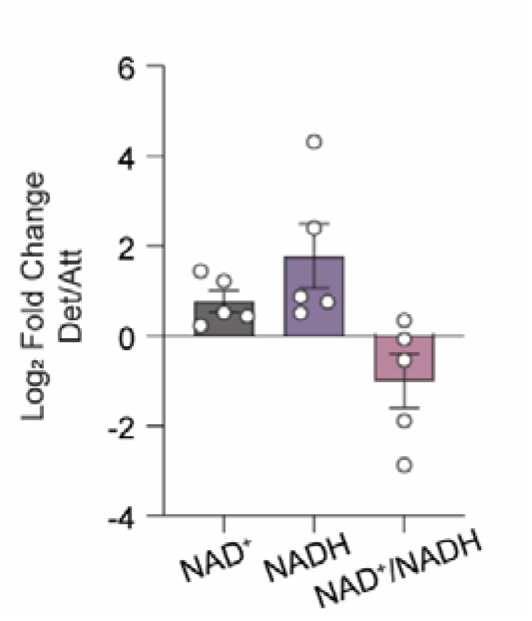
Trend toward NADH accumulation in detached compared to attached retina. Log_2_ fold change in detached versus attached retinal levels of NAD+ and NADH as well as the NAD^+^/NADH ratio after 24 hours. Two attached or detached retinas were pooled for each N with N=5 for both the attached and detached mouse groups. NAD^+^: nicotinamide adenine dinucleotide, NADH: nicotinamide adenine dinucleotide + hydrogen.

**Supp. Table 1.**
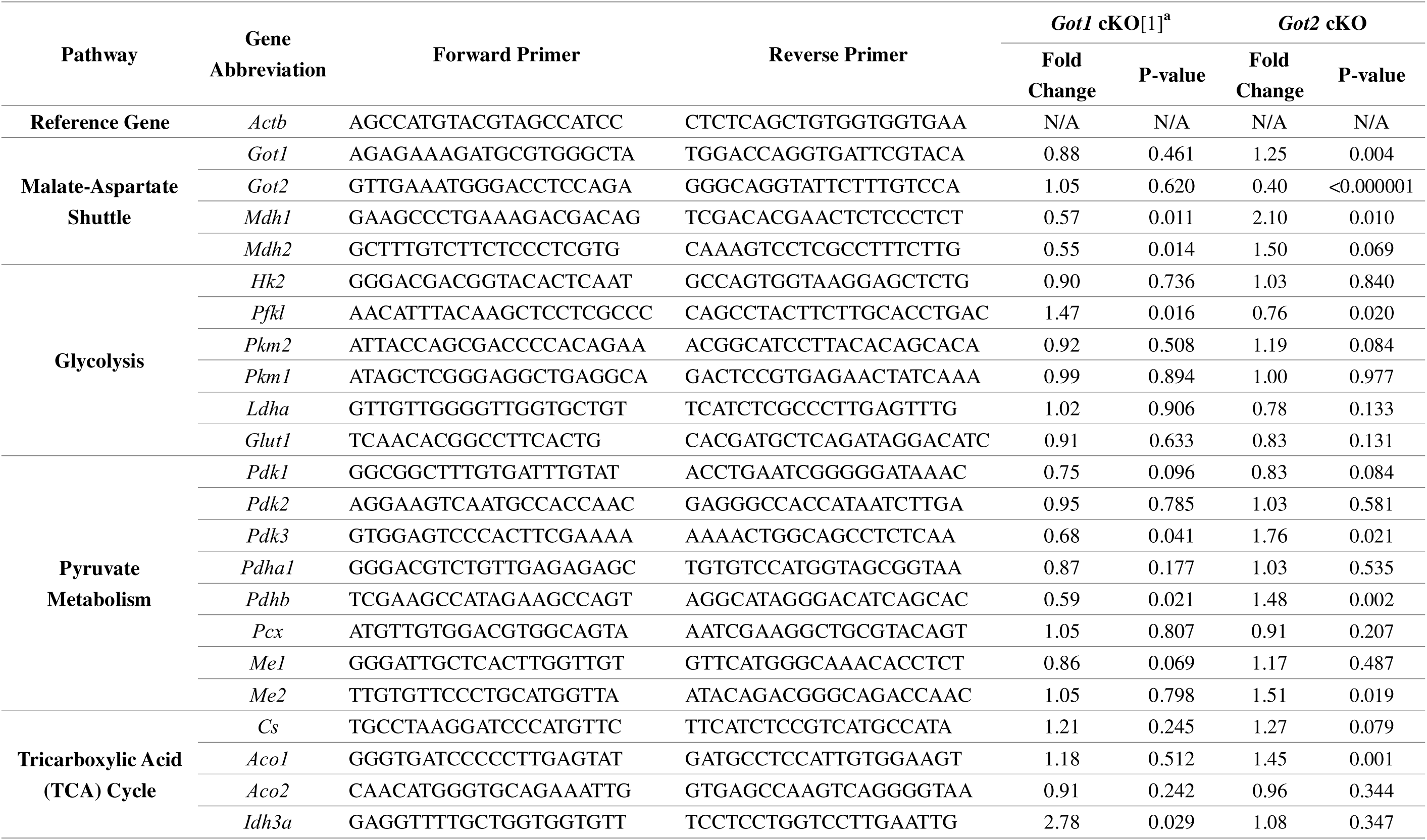
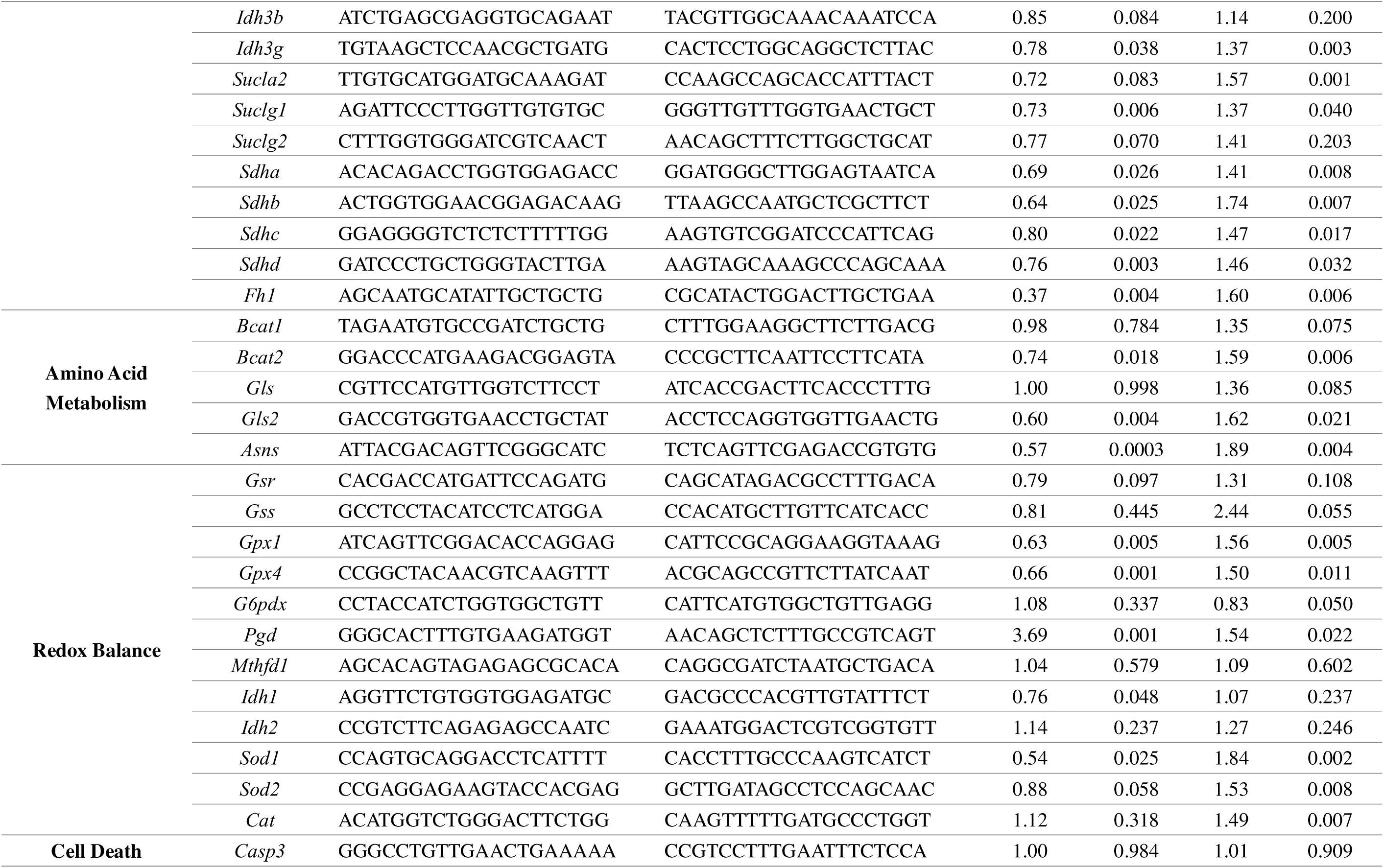
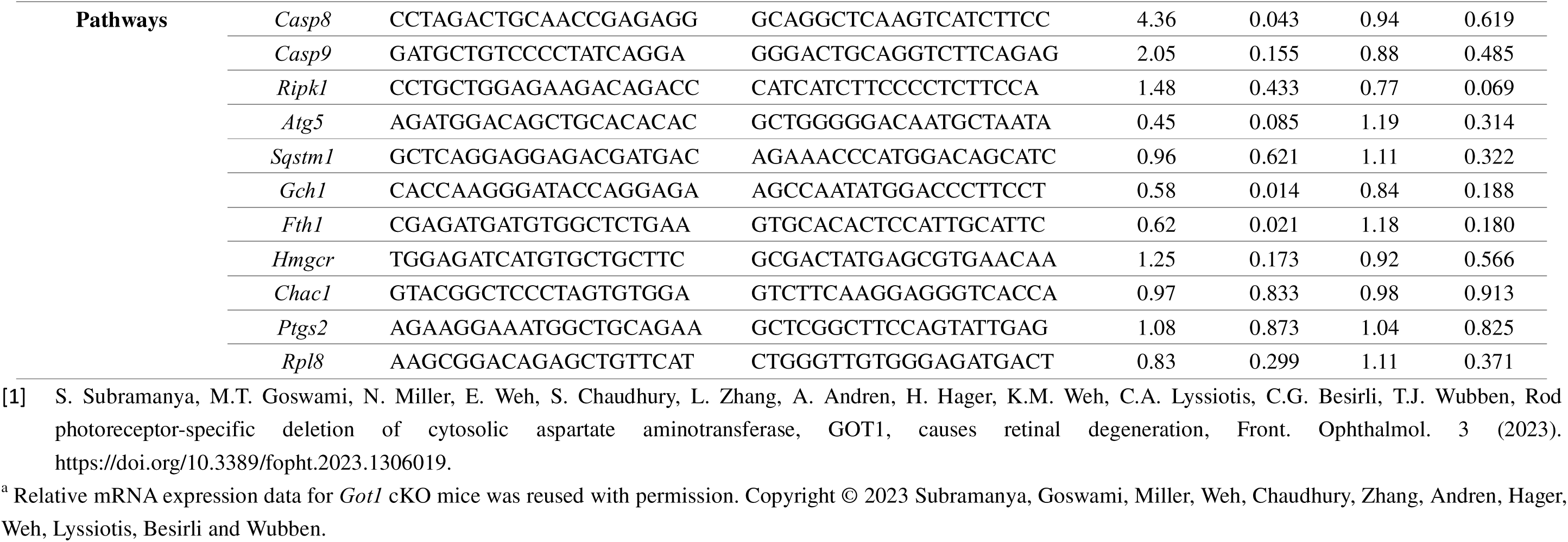
Gene expressions changes in *Got1* cKO and *Got2* cKO versus WT retina at 2 months of age.

